# Naphthalene monoimide derivative ameliorates amyloid burden and cognitive decline in a transgenic mouse model of Alzheimer’s disease

**DOI:** 10.1101/2020.08.20.260166

**Authors:** Sourav Samanta, Kolla Rajasekhar, Madhu Ramesh, N. Arul Murugan, Shadab Alam, Devanshi Shah, James P Clement, Thimmaiah Govindaraju

## Abstract

Alzheimer’s disease (AD) is a major neurodegenerative disorder and the leading cause of dementia worldwide. Predominantly, misfolding and aggregation of amyloid-β (Aβ) peptides associated with multifaceted toxicity is the neuropathological hallmark of AD pathogenesis and thus, primary therapeutic target to ameliorate neuronal toxicity and cognitive deficits. Herein, we report the design, synthesis and evaluation of small molecule inhibitors with naphthalene monoimide scaffold to ameliorate *in vitro* and *in vivo* amyloid induced neurotoxicity. The detailed studies established TGR63 as the lead candidate to rescue neuronal cells from amyloid toxicity. The *in silico* studies showed disruption of salt bridges and intermolecular hydrogen bonding interactions within Aβ42 fibrils by the interaction of TGR63, causing destabilization of Aβ42 assembly. Remarkably, TGR63 treatment showed a significant reduction in cortical and hippocampal amyloid burden in the progressive stages of APP/PS1 AD mice brain. Various behavioral tests demonstrated rescued cognitive deficits. The excellent biocompatibility, BBB permeability and therapeutic efficacy to reduce amyloid burden make TGR63 a promising candidate for the treatment of AD.

## INTRODUCTION

Alzheimer’s disease (AD) is a chronic neurodegenerative disorder contributing to 70-80% of all dementia cases (*1*). The phenotypic continuum describes the disease pathophysiology, parenchymal amyloid beta (Aβ) plaques deposition in the brain, cognitive decline, and neuropsychiatric symptoms viz., agitation, irritability, hallucinations and depression in the advanced stages (*2–5*). Aging is one of the risk factors, and more than 130 million people are expected to suffer from AD by 2050 (*6*). The complex disease etiology and lack of reliable treatments contributed to 146% rise in deaths by AD between 2000 and 2018 compared to an appreciable decline in the number of deaths caused by other disease conditions viz., heart disease, stroke, AIDS, prostate and breast cancer (*1, 6*). The National Institute on Aging and Alzheimer’s Association (NIA-AA) research framework report (2018) proposed parenchymal Aβ plaques as designated pathological hallmarks along with intracellular neurofibrillary tangles (*7*). The overexpression and proteolysis of amyloid precursor protein (APP) by β- and γ-secretases generate extracellular Aβ peptides which undergo misfolding and accumulate as senile plaques in the brain (*2, 5, 8, 9*). Specifically, Aβ42 aggregation species are incredibly neurotoxic and elicit toxicity in the form of disrupting neuronal synaptic function and plasticity, impaired short-term memory (STM), and long-term potentiation (LTP), a key process associated with learning and memory (*10, 11*). The endocytosis and blocking of essential receptors such as N-methyl-D-aspartate (NMDA) and α-amino-3-hydroxy-5-methyl-4-isoxazolepropionicacid (AMPA) at synaptic cleft by Aβ aggregation species cause neuronal circuit disruption and cognitive decline (*12–14*). The clinicopathologic studies correlate cognitive decline or sequence of AD pathology to Aβ burden associated toxicity in the AD brain (*8, 15, 16*). These facts and reports have underscored the importance of clearing or reducing the Aβ burden from the brain as a primary target to develop therapeutic agents for the treatment of AD (*17–19*). The amyloid toxicity emphasizes the need for a novel drug design strategy to ameliorate Aβ burden-associated plasma membrane toxicity, cognitive decline and memory (STM and LTP) impairment under progressive AD pathogenesis (*20–22*). We designed and synthesized a set of novel small molecules (TGR60-65) with naphthalene monoamide scaffold and evaluated their efficacy in ameliorating the amyloid toxicity of AD. The detailed biophysical, microscopy and cellular studies showed that 4-ethynyl-*N,N*-dimethylaniline and *N,N,N*-trimethylethylenediamine functionalized naphthalene monoimide (TGR63) are potent candidates to modulate Aβ42 aggregation and associated plasma membrane toxicity. The pharmacokinetics studies revealed serum stability, blood-brain-barrier (BBB) permeability and biocompatibility of TGR63 and its suitability for the long-term *in vivo* administration. The *in vivo* TGR63 treatment reduces the severe cortical and hippocampal Aβ burden in the APP/PS1 mice brain with significant improvement of memory and cognitive functions. Molecular dynamics study of Aβ species in the presence of TGR63 demonstrates the affinity and key interactions of TGR63 with Aβ peptides and provides insights on the modulation of toxic amyloidosis. *In vitro* and *in vivo* data on amelioration of amyloid burden, neuropathological hallmarks and cognitive decline establishes TGR63 as a potential therapeutic candidate to treat AD progression.

## RESULT

### Design and synthesis of small molecules with naphthalene monoimide scaffold

The Aβ aggregation causes deleterious neuropathological and cognitive effects and obliterating the amyloid burden and associated neurotoxicity are primary therapeutic strategies (*5, 22, 23*). We designed and synthesized a set of novel small molecules with naphthalene monoimide (NMI) scaffold to modulate the amyloid burden. The hydrophobicity of NMI core with *N,N,N*-trimethylethylenediamine as imide substituent was fine-tuned systematically by incorporating electron-rich *N,N*-dimethylamine, ethynylbenzene, and 4-ethynyl-*N,N*-dimethylaniline moieties at 8^th^ position (Fig. 1A, Scheme S1). These modifications to NMI core were undertaken to determine the required hydrophobicity-hydrophilicity to modulate Aβ aggregation. The structural fine-tuning of hydrophobic and hydrophilic moieties on NMI scaffold using appropriate functional groups resulted in a focused library of small molecules TGR60-65. For synthesis, 4-bromo-naphthalene monoanhydride (4-bromo NMA) was subjected to Sonogashira coupling with *N,N*-dimethylamine, ethynylbenzene, and 4-ethynyl-*N,N*-dimethylaniline using Pd(PPh_3_)_4_, sodium ascorbate and copper sulfate under argon atmosphere (Fig. 1A, Scheme S1). The NMA derivatives were conjugated with *N,N*-dimethylpropan-1-amine, *N,N,N*-trimethylpropan-1-aminium and 2-propoxyethan-1-ol in isopropanol under reflux (80 °C) conditions to obtain NMI derivatives TGR60-65 in good yields. All the final compounds were systematically characterized by nuclear magnetic resonance (NMR) and high-resolution mass spectrometry (HRMS).

**Fig. 1.**
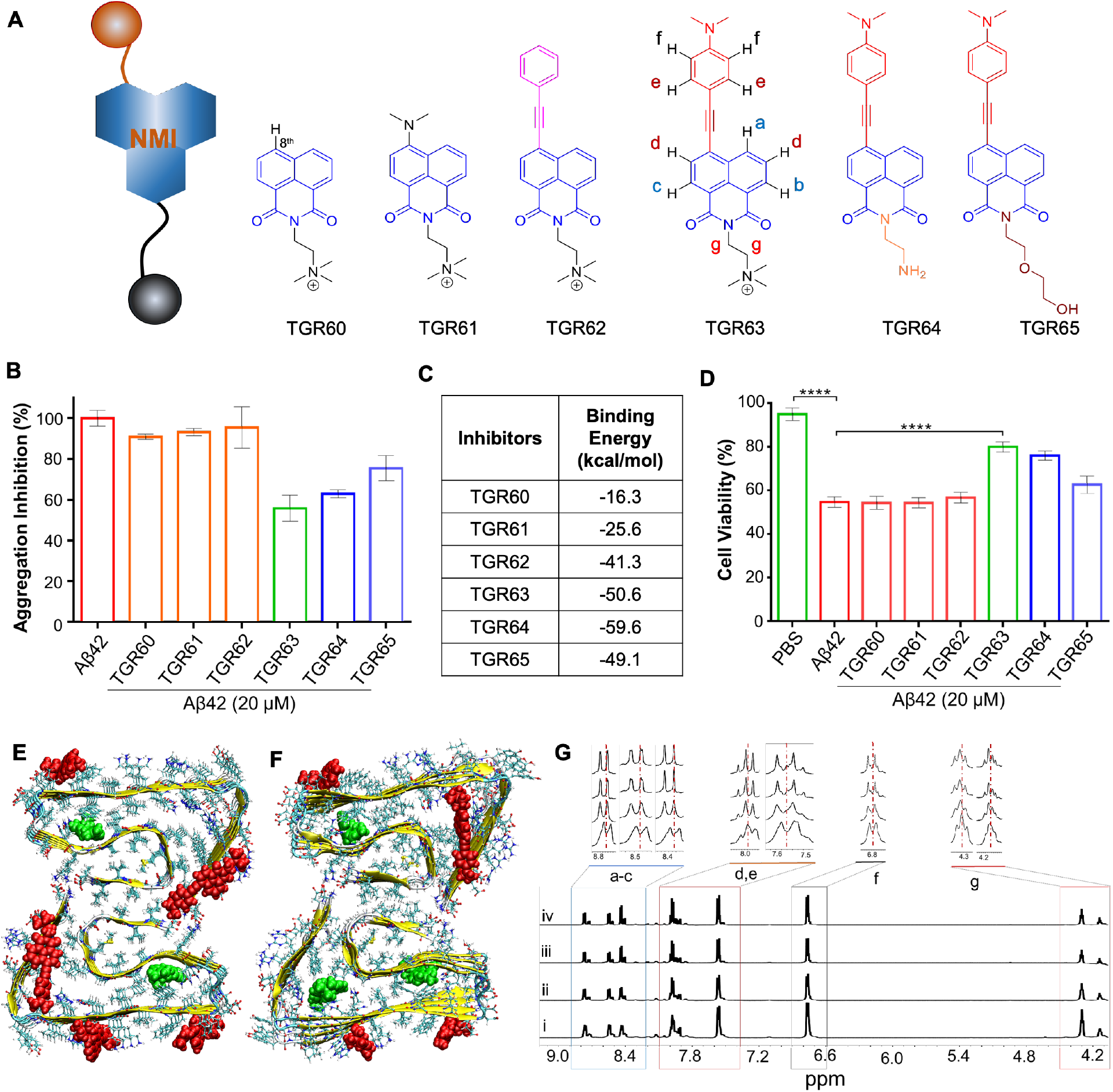
Design strategy of modulators of amyloid aggregation with NMI scaffold. (**A**) The core design of NMI derivatives and their chemical structures (TGR60-65). (**B**) Aggregation inhibition ability assessment: Aβ42 (10 μM) was incubated alone and with individual inhibitor (10 μM) for 72 h in PBS (10 mM, pH= 7.4) and the extent of aggregation was quantified by ThT fluorescence intensity. (**C**) Relative binding affinities of TGR60-65 compounds towards Aβ42 fibrils. The data are based on the top high affinity binding sites within fibrils. (**D**) *In vitro* neuronal rescue from amyloid toxicity by TGR60-65. The observed cell viability of cultured neuronal cells (PC12) after independently incubated with PBS, Aβ42 (20 μM) alone and in the presence of TGR60-65 (40 μM) for 24 h in cell growth media. (**E**) and (**F**) The high affinity binding sites of TGR63 within Aβ42 fibrils. E refers to the initial configuration, while F refers to a representative configuration in the production run. (**G**) Molecular level interactions between TGR63 and Aβ42: ^1^H NMR spectra of TGR63 in the absence (i) and presence of Aβ42 (10 μM) at 6 h (ii), 24 h (iii) and 48 h (iv) of incubation at 37 °C. Selected regions were magnified to show the change in proton chemical shift values due to molecular level interactions of TGR63 with Aβ, which are responsible for interference and disruption of noncovalent forces that drive Aβ aggregation. The chemical structure of TGR63 (A) with protons (a-g) assignment in different chemical environment.

### *In vitro* modulation of amyloid aggregation and probable mechanism

The ability of NMI derivatives (TGR60-65) to modulate Aβ42 aggregation and associated neuronal toxicity was evaluated through inhibition and dissolution assays. Thioflavin (ThT) fluorescence assay was employed to assess the aggregation modulation efficacy of TGR60-65. Aβ42 (10 μM) was incubated alone and with NMIs (10 μM) individually for 72 h, and the fluorescence intensity at 482 nm was measured upon treatment with ThT (10 μM) to assess the extent of aggregation (Fig. 1B). TGR60-62 treated Aβ42 samples showed 90%, 93% and 95% aggregation, respectively, compared to untreated control (100%), which corresponds to nominal inhibitory effects of ~10%, 7% and 5%, respectively. Aβ42 samples incubated with TGR63-65 showed ~55%, 62% and 75% aggregation suggesting significant aggregation inhibition of ~45%, 38% and 25%, respectively (Fig. 1B). Next, fully grown Aβ42 aggregates (10 μM) was incubated with TGR63-65 (30 μM) to assess their dissolution ability. The results showed a decrease in ThT fluorescence to 9%, 14% and 13%, which corresponds to ~91%, 86% and 87% dissolution of aggregates, respectively, in the presence of TGR63-65 compare to control (Fig. S1). These preliminary studies revealed that TGR63 is a promising lead aggregation modulator with pronounced inhibition and dissolution efficiency of 45% and 91%, respectively. A thorough computational study was carried out by employing molecular docking, molecular dynamics, and molecular mechanics-Generalized Born surface area (MM-GBSA) method to understand the molecular mechanism behind experimentally observed modulation of Aβ aggregation by TGR60-65 (*24, 25*). The Aβ42 fibril structure reported in the protein databank (pdb id is 5OQV) based on cryogenic-electron microscopy (cryo-EM) was used for this study. The study showed that molecules have the tendency to bind to multiple binding sites in Aβ42 fibril (Fig. 1E refers to binding sites for TGR63). The surface sites are shown in red color, and core sites are shown in green color. The binding free energies of TGR60-65 in their high-affinity binding sites were found to be −16.3, −25.6, −41.3, −50.6, −59.6, −49.1 kcal/mol, respectively (Fig. 1C). The lower binding free energies for TGR63-TGR65 indicate their better binding affinity and inhibition efficiency towards Aβ42 fibrils (Fig. 1B). These results encouraged us to assess the modulation of Aβ toxicity under cellular conditions (Fig. 1D). The AD-like environment was mimicked by exposing the cultured PC12 cells to Aβ42 (20 μM), which result in the generation of cytotoxic aggregation species in the growth media. Aβ42 caused mutilation to the cultured neuronal cells (PC12), as revealed by the decreased cell viability (54%) compared to untreated control cells (100%). The cells treated with Aβ42 in the presence of TGR60, TGR61, TGR62 and TGR65 showed cell viability of ~54%, 54%, 56% and 62%, respectively, similar to that of only Aβ treated cells. Interestingly, the promising aggregation modulators TGR63 and TGR64 showed 80% and 76% viability of cells treated with Aβ42, respectively (Fig. 1C). This rescue in viability corresponds to improved cell rescue of ~26% and 22% from Aβ toxicity by TGR63 and TGR64, respectively, with TGR63 displaying superior neuronal rescue effect. These findings were further confirmed by the cell rescue study using SHSY5Y cells, and the results are in good agreement with PC12 cells rescue data. TGR63 showed maximum cell rescue (~27%) effect on SHSY5Y cells compared to TGR64 (~3%) (Fig. S2). This neuronal rescue assessment confirmed that the 4-ethynyl-*N,N*-dimethylaniline and *N,N,N*-trimethylethylenediamine functionalization of NMI core (in TGR63) provides the best Aβ42 aggregation inhibition ability compare to other functional moieties. A concentration-dependent effect on the neuronal rescue was performed by treating SHSY5Y cells with varying concentrations of TGR63 (20, 40 and 100 μM) in the absence and presence of Aβ42 (20 μM) for 24 h. The data showed concentration-dependent cellular rescue with ~63%, 83% and 95% viability of cells observed for 20, 40 and 100 μM of TGR63, respectively, in the presence of Aβ42 (Fig. S3). In addition, the cytotoxicity assay of TGR63 in the absence of Aβ42 did not show any significant cytotoxicity at 100 μM compared to untreated control cells (100%), which demonstrates that TGR63 is nontoxic to cells at higher concentrations. *In silico* analysis was performed to understand the molecular mechanism behind the modulation of Aβ aggregation-induced toxicity by potential inhibitors (TGR63 and TGR64). Interestingly, both the inhibitors interact with Aβ42 fibril through multiple binding sites. In the course of molecular dynamics (MD), the inhibitors were found to bind to a “cryptic” site (a hidden site created when the ligand approaches the target) (Fig. 1E and F) (*24, 25*). The presence of such cryptic sites in Aβ42 fibril for novel inhibitor interaction is observed for the first time. The alterations in essential interactions (number of hydrogen bonds and salt bridges) of Aβ42 peptides that are mainly responsible for amyloid fibril formation due to the binding of TGR63 and TGR64 were analyzed. Aβ42 fibril comprised the maximum number of intermolecular hydrogen bonds (81 bonds) in the absence of inhibitor and was reduced to 75 and 74 in the presence of TGR63 and TGR64, respectively (Table S1). Further, TGR63 binding significantly reduced the salt bridges in Aβ42 fibrils from 48 to 41 through cation mediated disruption of electrostatic interactions (Fig. 1F and Table S1). However, certain new salt bridge interactions (total 54 interactions) were formed in the presence of TGR64 when compared to untreated Aβ42 fibrils. The observed changes in the hydrogen bonding and salt bridge interactions clearly explain the superior amyloid aggregation inhibition and dissolution potential of TGR63 compared to TGR64 (Table S1 and Fig. S4). A detailed analysis of the binding profile of TGR63 was performed due to its relatively superior disruptive interaction with Aβ42 fibril. There are two modes of binding for TGR63, i) core binding and ii) surface binding (Fig. 1F). The binding free energies of TGR63 in the core binding sites (as shown in green color in Fig. 1E) are associated with the least binding free energies (−50.6 & −47.8 kcal/mol), while that of the cryptic site is −34.9 kcal/mol (Fig. 1F). The TGR63 showed slightly higher binding free energies for the surface sites (−35.4, −30.3, −24.4, and −14.5, respectively) when compared to core sites (Table S2). The total binding free energies and individual contributions from the van der Waals, electrostatic, polar and non-polar solvation free energies are shown in Table S2. The data reveal that the Aβ fibril-TGR63 interaction is mostly driven by electrostatic and van der Waals interactions (−310.1 and −71.8 kcal/mol, respectively, for site-1). While the electrostatic interactions appear prominent, they are largely nullified by the polar solvation free energies making the van der Waals interactions as the major driving force for the ligand-fibril association. The *in silico* study revealed that the binding of TGR63 at the surface and core sites of fibrils is responsible for modulation of Aβ aggregation. Overall, the *in vitro, in cellulo* and *in silico* studies, established TGR63 as a promising candidate to modulate Aβ aggregation and associated toxicity in cells.

Next, NMR spectroscopy was used to ascertain the molecular-level interactions between TGR63 and Aβ42 peptide (Fig. 1G). ^1^H NMR spectra of TGR63 (1 mM) were acquired in the absence and presence of Aβ42 (10 μM) at different incubation time points (24, 48 and 72 h) using WATERGATE sequence for solvent suppression in PBS buffer (10 mM, pH= 7.4) containing D_2_O (12%) (*20*). TGR63 alone showed aromatic protons of NMI core and aniline moiety (a-f) in the chemical shift range of 6.5–8.8 ppm (Fig. 1G). In the presence of Aβ42, the splitting pattern of these aromatic protons (7.4–8.8 ppm) was completely altered with a significant downfield shift (~0.05 ppm) as a function of time. This reorganization of NMR signals confirmed the interactions between the aromatic moieties of TGR63 and Aβ42 peptide and are responsible for the observed aggregation modulation. In addition, the ethyl protons (g) signals at 4.1–4.4 ppm became sharper with time and experienced a significant downfield shift, which indicates the interactions of ethyl protons of TGR63 with Aβ42 peptide. As discussed (*vide supra*), the aggregation-prone Aβ42 peptides readily self-assembles into the ordered β-sheet structure through noncovalent interactions (*2, 5, 8*). The NMR data provided insights into the molecular level interactions of TGR63 with Aβ42 that possibly modulate the Aβ42 aggregation by disruption of crucial noncovalent interactions.

### Inhibition of Aβ aggregation and associated toxicity: microscopy and dot blot analysis

Modulation of Aβ aggregation by TGR63 was evaluated through the structural and morphological analysis using atomic force microscopy (AFM) and transmission electron microscopy (TEM). Aβ42 (10 μM) was incubated alone and with TGR63 (50 μM) for 48 h in PBS (10 mM, pH= 7.4), and the samples were spotted on a mica surface and TEM grid to acquire AFM and TEM images, respectively (Fig. 2A and B). AFM image of Aβ42 sample showed long fibrillar structures with ~3.0 nm height, while TGR63 treated Aβ42 sample revealed amorphous structures (Fig. 2A). Similarly, TEM image displayed a highly intertwined fibrillar structure of Aβ42, which is significantly disrupted by the treatment with TGR63 (Fig. 2B). The modulation (inhibition and dissolution) of Aβ aggregation was further supported by the dot blot (immunoassay) analysis (Fig. 2C). Aβ42 (10 μM) samples were incubated alone or with different concentrations of TGR63 (10 and 50 μM) for 48 h at 37 °C. The incubated samples were spotted on a polyvinylidene difluoride (PVDF) membrane and probed with Aβ fibrils specific OC (1:1000) primary antibody followed by secondary antibody (1:10000). The spots on the PVDF membrane were further treated with enhanced chemiluminescence (ECL) reagent to image and assess the extent of inhibition of Aβ aggregation using a Versa Doc instrument. The blot image and their quantification data revealed the maximum amount of fibrillar aggregates for Aβ42 (L1) sample (100%). In comparison, a significant reduction of fibrillar aggregates was observed in the presence of TGR63 (10% and 60% for L2: 10 μM and L3: 50 μM, respectively) in a concentration-dependent manner (Fig. 2C). These results and observations from AFM, TEM and dot blot analysis have validated the data from ThT fluorescence assay to confirm TGR63 as a potential modulator of Aβ aggregation.

**Fig. 2.**
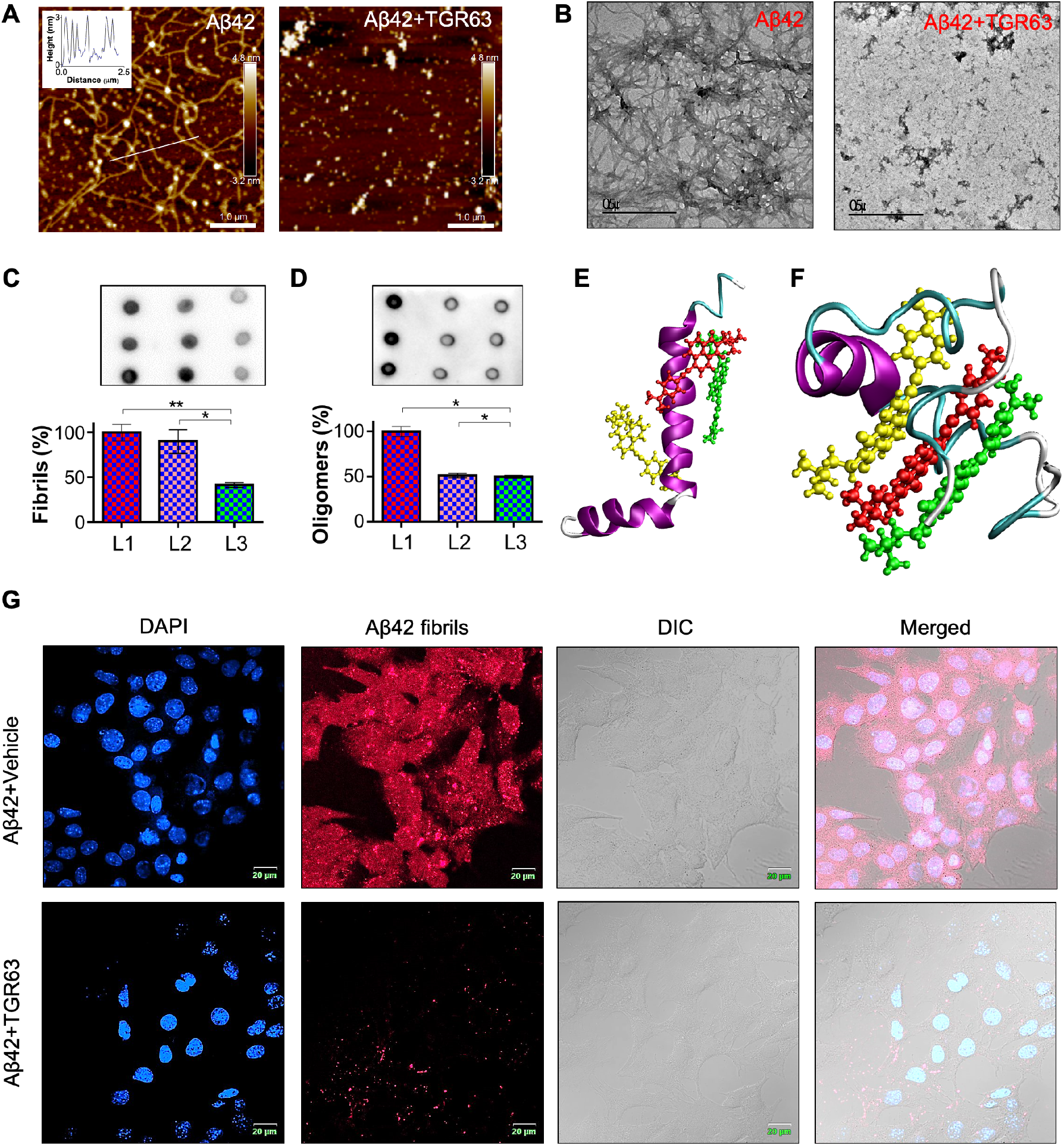
Modulation of Aβ aggregation and associated toxicity. (**A**) AFM images of Aβ42 in the absence (inset: height profile) and presence of TGR63. (**B**) TEM images of Aβ42 in the absence and presence of TGR63. (**C**) Dot blot analysis of TGR63 treated Aβ42 fibrils: The blot intensity displayed the amount of Aβ42 fibrils (5 μM) in absence (L1) and presence of TGR63 at two different molar ratios 1:1 (L2) and 1:5 (L3). Aβ42 fibrils were probed using OC primary antibody (1:1000) and treated with ECL reagent to capture the image in Versa Doc instrument and the comparison of blot intensities (%) revealed the effect of TGR63 in amyloidosis. (**D**) Dot blot analysis of TGR63 treated Aβ42 oligomers: The blot intensity displayed the amount of Aβ42 oligomers (5 μM) in the absence (L1) and presence of TGR63 at two different molar ratios 1:1 (L2) and 1:5 (L3). Aβ42 oligomers were probed using A11 primary antibody (1:1000) and treated with ECL reagent to capture the blot image and the comparison of blot intensities (%) revealed the inhibition effect of TGR63 in Aβ42 oligomerization. (**E**) The most stable (least energy) binding modes for TGR63 in monomeric Aβ42 peptide. E also refers to input MD configuration. (**F**) refers to a representative configuration for the TGR63-monomeric Aβ42 peptide complex during the production run. (**G**) Protection of plasma membrane from Aβ toxicity: Confocal microscopy images of SHSY5Y cells after incubating (2 h) independently with only Aβ42 (10 μM) fibrils (Aβ42+Vehicel) and TGR63 (50 μM) treated Aβ42 (10 μM) fibrils (Aβ42+TGR63). The SHSY5Y cells were stained with OC (1:250) primary antibody followed by fluorescently (λ_ex_= 633 nm, λ_em_= 650 nm) labeled (red) secondary antibody (1:250) and DAPI (blue). Scale bar: 0.5 μm (TEM); 1.0 μm (AFM); 20 μm (confocal).

The soluble Aβ aggregation species, namely oligomers, are considered highly toxic and critical contributors to Aβ toxicity (*5, 8*). Aβ oligomers are known to interact and disrupt the lipid membranes (mitochondrial and plasma membranes), causing mitochondrial dysfunction and neuronal damage (*18*). The disruption of the plasma membrane at synaptic cleft causes synaptic dysfunction, followed by weakening in synaptic plasticity and LTP formation (*10–14*). Therefore, a potential modulator of Aβ aggregation must effectively inhibit oligomer formation to protect neuronal cells and improve memory impairment in AD. We assessed the inhibitory activity of TGR63 against Aβ42 oligomers by immunohistochemistry assay (Fig. 2D). Aβ42 monomers (10 μM) were incubated in the absence (L1) and the presence of varying concentrations of TGR63 (10 and 50 μM) for 24 h at 4 °C. The incubated samples were spotted on the PVDF membrane and treated with Aβ oligomer-specific primary antibody (A11) followed by ECL reagent to image and quantify the extent of inhibition of Aβ oligomers using Versa Doc instrument (*26*). The quantification of spot intensities showed ~48% and 50% inhibition of oligomer by TGR63 (10 and 50 μM, respectively) treatment compared to untreated control (100%) (Fig. 2D). *In silico* analysis was performed using an integrated approach (molecular docking, molecular dynamics and binding free energy calculations) to understand the effect of TGR63 on the conformational dynamics of monomeric Aβ42 peptides. The α-helix structure of Aβ42 is essential for the formation of oligomers and their interaction to disrupt the lipid membrane structure. TGR63 induced secondary structural changes to play a key role in the oligomer formation kinetics and membrane toxicity. The molecular docking showed three different low energy binding modes (site 1m-3m) for TGR63 in monomeric Aβ peptide (Fig. 2E and F). The binding free energies in 1m-3m sites were found to be −24.1, −9.4 and −26.5 kcal/mol, respectively (Table S3). It is worth noting that the binding free energies for TGR63 with monomeric Aβ peptide are higher compared to binding with fibril (Table S2). The considerable reduction in the van der Waals interactions in case of the former contributes to observed differences in the binding energies. Figures 2E shows Aβ peptide (with α-helix contents ~76%) structure similar to the fusion domain of virus influenza hemagglutinin, which is responsible for making holes and causes plasma membrane damage (*24*). Interestingly, TGR63 treatment effectively reduced the α-helix content of Aβ peptide, which resulted in the formation of a nontoxic globular structure (Fig. 2F). Overall, the blot analysis and *in silico* assessments validated that TGR63 is an efficient modulator of polymorphic species of Aβ aggregation and a potential candidate to ameliorate the amyloid burden and associated membrane toxicity.

Membrane toxicity induced Aβ aggregation species is one of the major toxicity routes to neuronal death (*5, 8, 27*). The deposition of Aβ plaques on the healthy axon and dendrons of mature neurons is a primary cause of neuronal atrophy in the AD brain (*13, 14*). In addition, soluble Aβ oligomers dampen smooth neuronal signaling by blocking the neuronal surface receptors (NMDA and AMPA) at synaptic cleft (*10, 12*). The synaptic dysfunction impairs the synaptic plasticity and hippocampal LTP formation causing neuronal damage, memory loss and cognitive decline under AD pathogenesis (*4*). Contemporary studies have shown that Aβ aggregation species interact with the plasma membrane and promote the internalization of misfolded Aβ peptides by punch holes through the membrane (5, *28, 29*). Inhibition of Aβ-membrane interaction and associated toxicity is anticipated to rescue neuronal cells from the amyloid burden. The protective effect of TGR63 on neuronal cells from the membrane toxicity caused by Aβ was evaluated in SHSY5Y cells using immunocytochemistry protocols. The cells were cultured in 35 mm confocal dishes and treated independently with Aβ42 and pre-incubated (24 h) Aβ42-TGR63 for 2 h in the cell growth media. The experimental cells were washed and fixed using 4% paraformaldehyde (PFA) and treated with OC (1:250) antibody, followed by red fluorescent-labeled (λ_ex_= 633 nm and λ_em_= 650 nm) secondary antibody. The unbound antibody was washed, and the cells were treated with nuclear staining dye DAPI for confocal imaging. The red fluorescence signal was significantly high and mostly localized on the plasma membrane for cells treated with Aβ42 (Aβ42+Vehicle), which correlates to levels of Aβ42 fibrillar aggregates (Fig. 2G). The cells treated with Aβ42-TGR63 sample showed a significant reduction in the red fluorescence signals on the plasma membrane. This observation supports the inhibition of toxic Aβ aggregation species by TGR63 to protect the plasma membrane. Collectively, the *in vitro* and *in cellulo* results showed the importance of simple structure-function relationship study and the balanced hydrophobicity and hydrophilicity of TGR63 attained by means of meticulously chosen substituents (4-ethynyl-*N,N*-dimethylaniline and *N,N,N*-trimethylethylenediamine) to modulate the Aβ aggregation as per the design strategy. These results motivated us to evaluate the anti-AD properties of TGR63 in an APP/PS1 double transgenic AD mouse model.

### Pharmacokinetics study of TGR63

We performed pharmacokinetics of TGR63 in Wild-type (WT) mice to assess it’s *in vivo* efficacy (Fig. 3A). The lethal dose 50 (LD50) of TGR63 was determined in WT mice through intraperitoneal (IP) injection following the Organisation for Economic Co-operation and Development (OECD) guidelines. Twenty-five WT mice were segregated in five different groups (G1-5, N= 5 per group) and administered with varying doses of TGR63 (1.7, 5.5, 17.5, 56.0 and 179.0 mg/kg body weight, respectively) through IP injection and their survival was monitored for 14 days (Fig. S5A). The survival of the experimental mice showed that TGR63 is mostly nontoxic in the experimental period due to the high LD50 value of ~157.9 mg/kg body weight (Fig. S5B). The serum stability and blood-brain barrier (BBB) crossing ability of TGR63 were assessed through matrix-assisted laser desorption ionization (MALDI) mass spectrometry analysis of blood and brain samples of vehicle and TGR63 treated mice. TGR63 and vehicle were administrated in WT mice and sacrificed after 1 and 24 h to collect the blood for MALDI mass analysis (Fig. 3B and Fig. S6). The MALDI analysis confirmed the presence of TGR63 in blood after 24 h of administration. TGR63 was incubated in PBS (10 mM, pH= 7.4) and blood serum in WT mice for different time intervals (0.5, 1, 2 and 6 h) at 37 °C to evaluate the serum stability under *in vitro* conditions. The spectrometric analysis (absorbance) confirmed the stability of TGR63 in blood serum (Fig. S7). Next, we calculated partition coefficient (P), a valuable physical property to predict the BBB permeability. TGR63 (20 μM) was added to an immiscible solution of water (10 mL) and octanol (10 mL), followed by thorough mixing, the solution was allowed to segregate into two layers. The absorption of the octanol layer at 450 nm was measured, and the amount of TGR63 was determined from the standard concentration curve (Fig. S8). The concentrations of TGR63 in octanol and water layer were found to be 10.45, and 29.55 μM, respectively and logP value was calculated to be 0.1 (Fig. S8). The calculated positive logP value predicts the possible BBB crossing ability for TGR63 (*18*). For *in vivo* assessment, TGR63 and vehicle administrated WT mice were sacrificed after 1 h to collect the brains for MALDI mass analysis. TGR63 treated mouse brain sample showed a mass peak at 426.04 (m/z), which was absent in the vehicle-treated sample and confirmed BBB crossing ability of TGR63 (Fig. 3C and Fig. S9). Further, TGR63 (5 mg/kg body weight) and vehicle (control) were administrated in age (6 months old) matched APP/PS1 and WT mice daily for 8 months to examine the organ toxicity upon prolonged TGR63 administration (Fig. 3D). The experimental mice were sacrificed at 14 months of age and critical organs *viz*., liver, heart, spleen and kidney were harvested to perform gold standard hematoxylin and eosin (H&E) staining (stain nucleus and cytoplasm, respectively). The H&E staining of TGR63 treated mice (WT and AD) tissue samples exhibited nucleus and cytoplasm staining similar to healthy tissue (vehicle-treated WT mice). The healthy or TGR63 treated tissue samples did not show any abnormal scar, disorganization, inflammatory infiltrate, hepatotoxicity or necrosis (Fig. S10), which confirmed the tremendous *in vivo* biocompatibility and nontoxic nature of TGR63. The pharmacokinetics study of TGR63 revealed serum stability, BBB permeability and biocompatibility, underscoring its suitability for the long-term treatment in APP/PS1 AD phenotypic mice. These studies have encouraged us to evaluate the efficacy of the lead candidate to ameliorate the cognitive impairment, for which APP/PS1 AD and WT mice were administrated (IP) with TGR63 (daily dose of 5 mg/kg body weight) starting from the age of 6 months to 14 months (Fig. 4D).

**Fig. 3.**
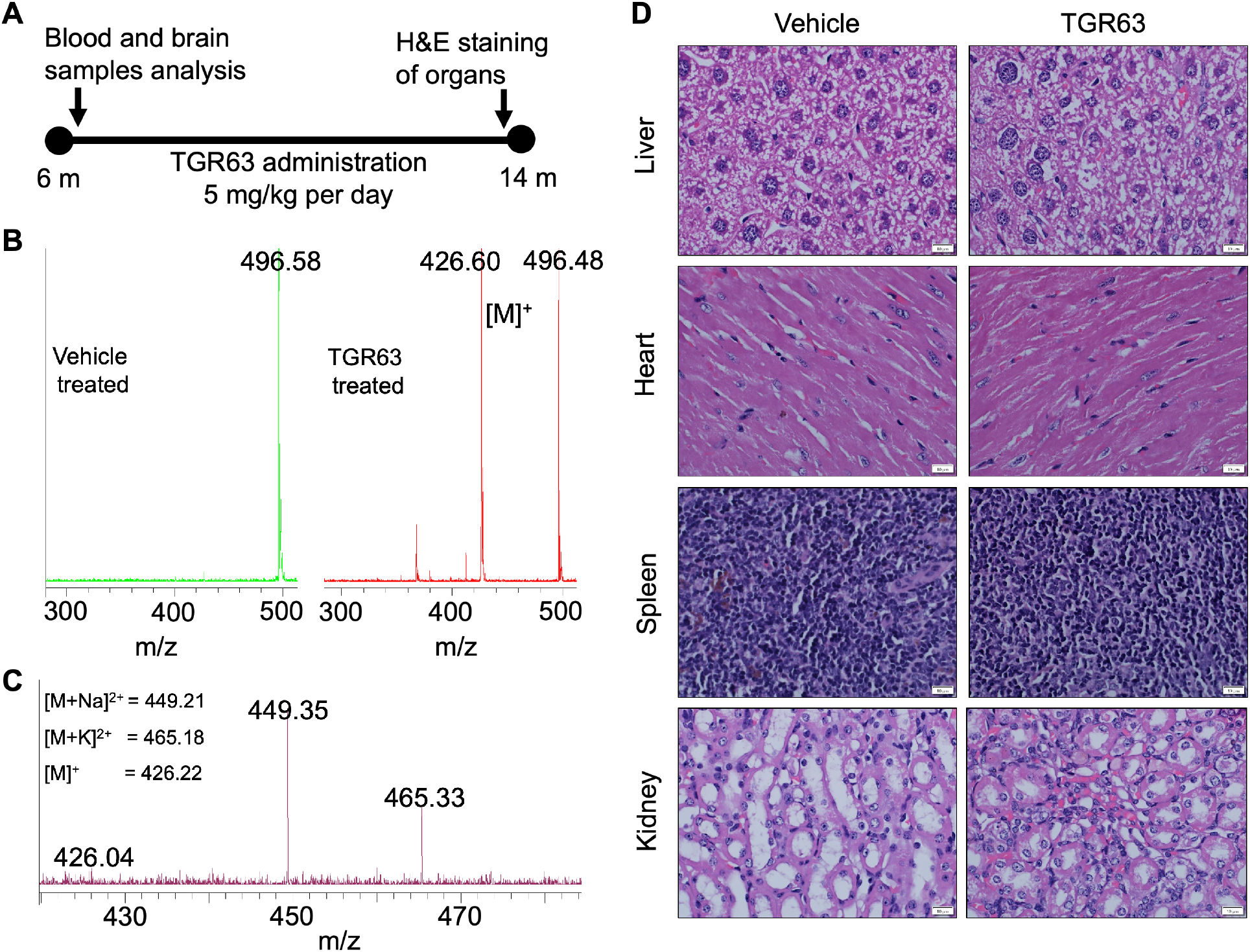
The *in vivo* blood brain barrier (BBB) permeability and pharmacokinetics study of TGR63. (**A**) Experimental planning and TGR63 administration in WT mice (age in month, m) to assess the pharmacokinetics. (**B**) Serum stability of TGR63: MALDI mass analysis of TGR63 treated mouse blood serum to detect the presence of TGR63 (calculated [M]^+^ = 426.21) in the blood after 1 h of administration and comparison with vehicle treated blood sample. (**C**) BBB crossing ability: MALDI mass analysis of TGR63 administrated mouse brain lysate, which showed the presence of TGR63 in the brain and established its BBB permeable ability. (**D**) Evaluation of organ toxicity of TGR63: Bright field images of vehicle and TGR63 treated WT mouse organs (liver, heart, spleen and kidney) stained with hematoxylin and eosin (H&E), which confirmed the biocompatibility and nontoxic nature of TGR63. Scale bar: 10 μm.

**Fig. 4.**
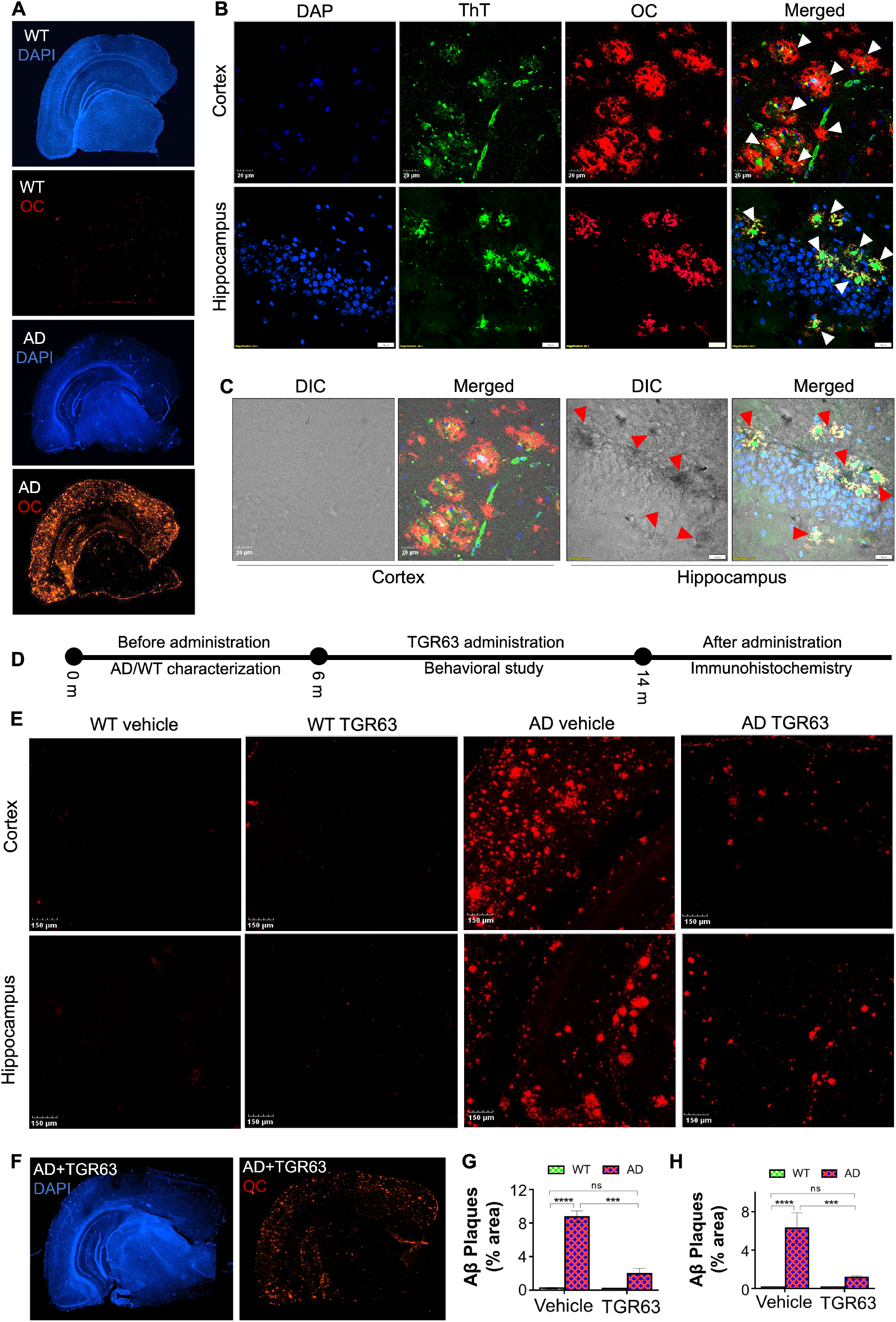
Reduction of amyloid burden by TGR63 in APP/PS1 AD phenotypic mice model. (**A**) Visualization of amyloid plaques in half hemisphere: Confocal microscopy images of coronal section of WT and AD mice brains immunostained with amyloid fibrils specific OC primary antibody followed by fluorescently (λ_ex_= 633 nm, λ_em_= 650 nm) labeled (red) and DAPI (blue) (**B**) Staining of amyloid plaques with OC primary antibody and ThT or CQ probe: The high-resolution confocal microscopy images of cortex and hippocampus regions of the AD mouse brain, immunostained with OC antibody (red), DAPI (blue) and ThT (green). The merged images display significant overlap between ThT and OC staining to confirm the amyloid deposition (pointed with white arrows). (**C**) Visualization of amyloid deposits associated neuronal damage: The DIC images of different regions of AD. The merged images of DIC and confocal microscopy images show amyloid plaques associated brain damage (pointed out with red arrows). (**D**) Experimental planning and TGR63 administration in APP/PS1 mice (age in month, m). (**E**) Reduction of cortical and hippocampal amyloid burden by TGR63 treatment: Higher magnification images of vehicle and TGR63 treated mice (WT and AD) brain sections to visualize and compare the Aβ plaques deposition in the cortex and hippocampus areas. The brain tissue sections were immunostained with amyloid fibrils specific primary antibody (OC) and red fluorescent-labeled (λ_ex_= 633 nm and λ_em_= 650 nm) secondary antibody. (**F**) Visualization of amyloid plaques in half hemisphere: Confocal microscopy images of coronal section of TGR63 treated AD mouse brain immunostained with amyloid fibrils specific OC primary antibody followed by fluorescently (λ_ex_= 633 nm, λ_em_= 650 nm) labeled (red) secondary antibody and DAPI (blue). (**G** and **H**) Quantification of Aβ plaques: Amount of Aβ plaques (% area) deposited in different regions (cortex and hippocampus) of vehicle and TGR63 treated mice (WT and AD) brain was analyzed. Data represent mean ± SEM, n= 3 per group. Scale bar: 20 μm.

### *In vivo* amelioration of amyloid burden

We sought to evaluate the activity of TGR63 to ameliorate amyloid burden in *in vivo* AD model. APP/PS1 mice were bred, maintained and characterized (WT: wild type; AD: APP/PS1 positive) according to the Jackson Laboratory protocols (*16*). The double transgenic APP/PS1 mice (B6C3-Tg (APPswe, PSEN 1dE9)85Dbo/J; stock number 004462), which express human transgenes APP and presenilin 1 (PS1) in the central nervous system (CNS) contains the Swedish and L166P mutations, respectively (*30*). The K595N/M596L (Swedish) mutation favors the amyloidogenic processing of APP protein, and PS1 mutation (L166P) elevates the production of Aβ peptides through modifying the intra-membrane γ-complex. Consequently, deposition of Aβ plaque starts appearing in the neocortex at the age of ~45 days and can be found in thalamus, brainstem, striatum and hippocampus regions at the age of 5-6 months. The deposition of Aβ plaque in the cortex and hippocampus regions initiate cognitive dysfunction and memory impairment at the age of ~7 months. The presence of Aβ plaques in the APP/PS1 AD phenotypic mouse brain was confirmed and compared with the healthy brain by Aβ plaques-specific staining protocols (Fig. 4A). The brains were harvested from the age matched WT and AD mice and treated with PFA (4%) and sucrose solution (30%) for the sagittal brain sectioning (40 μm sections). The brain sections were co-stained with ThT (λ_ex_= 442 nm, λ_em_= 482 nm) and OC primary antibody followed by fluorescently labeled secondary antibody (λ_ex_= 633 nm, λ_em_= 650 nm) or CQ to visualize and confirm the amyloid plaques deposition (*37*). The confocal images acquired from different regions of the brain (cortex and hippocampus) showed localized bright green and red fluorescence signals confirming the deposits of amyloid plaques in the APP/PS1 mice brain. Similar fluorescence signals (green and red) were absent in the age-matched WT brain section, confirming the amyloid plaques-free healthy brain (Fig. 4B). The hippocampal damage, a hallmark of advanced AD condition was partially observed in 14 month old APP/PS1 mice (Fig. 4C). Age-matched AD, and WT cohorts were administered with TGR63 (5 mg/kg body weight/day) and vehicle starting from the age of 6 months following our treatment protocols (Fig. 4D). The experimental mice were sacrificed after completing the behavioral studies (14 months) to investigate amyloid deposits in the brain using immunohistochemistry (*16*). The sagittal brain sections were permeabilized and blocked with PBTx (0.1M PBS and 0.1% TritonX-100) and goat serum (1%) containing BSA (2%) at room temperature, respectively. The processed sections were incubated with amyloid fibrils specific primary antibody (OC, 1:250) at 4 °C for 48 h to stain the dense core of amyloid plaques. The processed brain sections were further treated with red fluorescent-labeled (λ_ex_= 633 nm and λ_em_= 650 nm) secondary antibody (1:1000) and DAPI to perform confocal imaging (Fig. 4E). The confocal images of WT cohort brain tissue sections did not show any deposits of Aβ plaques in both cortex and hippocampus regions. The age-matched AD cohort brain tissue sections prominently displayed deposits of Aβ plaques in different parts of the brain *viz*., neocortex, striatum, primary sensory-motor areas, hippocampus, temporobasal and frontomedial areas (Fig. 4F). These results provided strong evidence of chronic accumulation of Aβ plaques in the brain associated with AD progression. Predictably, the vehicle-treated AD brain tissue images (N= 3) showed an accumulation of Aβ plaques 8.87% and 6.28% area of the cortex and hippocampus, respectively (Fig. 4E). Remarkably, TGR63 treatment (N= 3) significantly reduced the Aβ plaques deposits to 1.94% and 0.94% area of the cortex and hippocampus, respectively (Fig. 4G and H). In other words, TGR63 treatment reduced Aβ deposits by 78% and 85% in the cortex and hippocampus, respectively. The immunostaining of Aβ deposits in TGR63 treated AD brain tissue displayed a considerable reduction in the amyloid load and encouraged us to test for the corresponding improvement of memory and cognitive functions.

### Recovery of cognitive functions

AD is characterized by the progressive deterioration in cognitive functions, which generally include learning and memory impairment leading to neuropsychiatric symptoms viz., aggression, agitation, anxiety and depression (*2, 4, 8*). APP/PS1 mice show age-related AD-like phenotypes linked to Aβ plaques deposition in the brain (*32*). We set out to assess the recovery of cognitive functions in TGR63 treated APP/PS1 mice (Fig. 5). Open-field (OF) test was performed to evaluate the effect of TGR63 on anxiety and locomotion. Next, the amelioration of learning disability and memory impairment by TGR63 treatment was evaluated through novel object identification (NOI) and Morris water maze (MWM) behavioral tests. In OF test, all the experimental mice were individually allowed to explore a novel platform (45 ≥ 45 cm) and their locomotion activity was monitored for 5 min (Sony HDRCX405 camera) and analyzed using the smart 3 software (Panlab; Fig. 5A, Fig. S11-14). The trajectories of vehicle-treated AD mice (AD vehicle) showed higher activity (travel average 2698.25 cm) compared to vehicle-treated WT mice (travel average 1533.88 cm), which indicates the AD-like phenotype of APP/PS1 mouse model (Fig. 5B). Interestingly, TGR63 treated AD (AD TGR63) mice showed significantly shorter travel paths (average 1515.33 cm) compared to AD vehicle cohort suggesting improved locomotor functions and anxiety similar to vehicle-treated WT mice (WT vehicle). The anxiety behaviors of TGR63 treated mice were assessed by the time spent and the entries in the center zone (20 ≥ 20 cm) of OF arena. As expected, AD vehicle showed the maximum number of entries (~20) and travel path (average 243.0 cm) among other cohorts in the center zone, which confirmed the characteristic anxious nature of AD conditions (Fig. 5C and D). Remarkably, TGR63 treated AD mice showed behaviors similar to healthy WT vehicle cohorts with ~9 entries and travel average of 98.14 cm exploration in the center zone. The OF test data revealed that TGR63 ameliorated the β-amyloid-induced aggression, agitation and anxiety observed in the middle stages of AD. Next, the effect of TGR63 on memory processing, viz., acquisition, consolidation and retrieval were evaluated through NOI test, which has been widely used as a tool to study the neurobiology of memory using the natural tendency of rodents to explore novel objects more than the familiar objects (*33*). All the experimental mice were familiarized with two identical objects (familiar objects) in a known habituated arena and allowed to explore a novel and familiar object after 24 and 48 h of familiarization (Fig. 5E). The exploration time with each object was measured using a stopwatch, and the discrimination index (DI) was determined using the formula, (time exploring the novel object - time exploring the familiar) / (time exploring novel + familiar) * 100 (*27*). The test result after 24 h showed significantly lower DI (−3) for AD vehicle cohort compared to WT vehicle cohort (+49), which affirmed the deteriorating memory formation and recollection under progressive AD conditions (Fig. 5F). On the other hand, calculated DI of WT TGR63 cohort (+50) is similar to the WT vehicle cohort confirming TGR63 did not affect memory formation. Remarkably, AD TGR63 cohort exhibited an improved DI (+43) compared to AD vehicle cohort (−3) confirming the therapeutic efficacy of TGR63 in memory processing (acquisition, consolidation and retrieval) under AD condition. Similarly, the calculated DI after 48 h was lowest (−7) for AD vehicle cohort compared to both vehicle and TGR63 treated WT cohorts (+43 and +45, respectively) (Fig. 5G). AD TGR63 cohort showed DI of +38, which indicate normal memory formation and retrieval. The DI of TGR63 treated WT, and AD cohorts at 48 h have marginally reduced (~ 5 units of DI) compare to 24 h, reveal the natural long-term depression of healthy animals. The blocking of essential synaptic receptors (NMDA and AMPA) by Aβ aggregation species leading to synaptic dysfunction followed by impairment in hippocampal LTP formation. The NOI test result demonstrated that AD positive mice (APP/PS1) exhibit the memory impaired phenotypes compare to WT mice. TGR63 treatment ameliorates the memory impairment in APP/PS1 mice by reducing the toxic amyloid burden from the brain under progressive AD conditions.

**Fig. 5.**
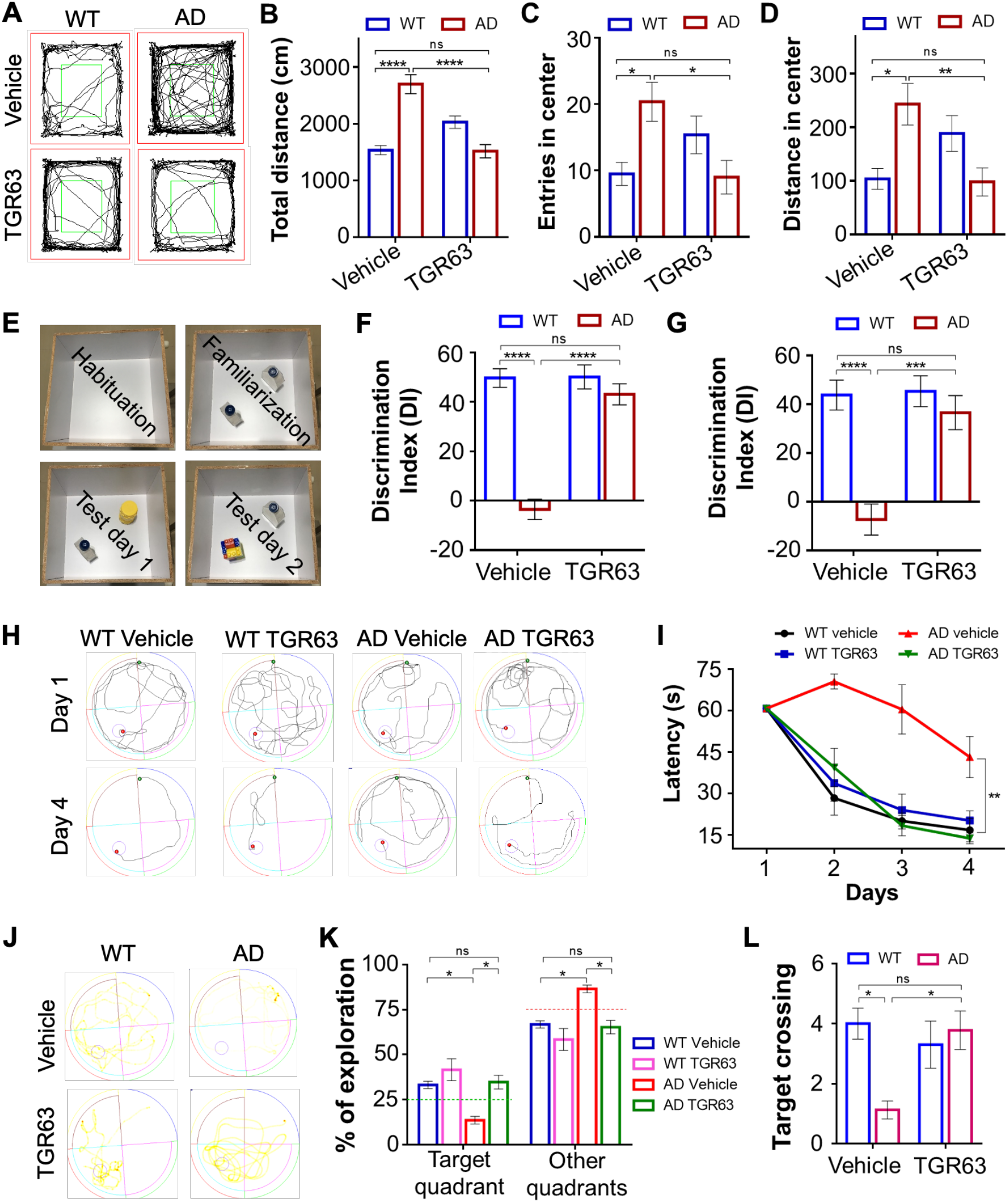
Improvement of memory and cognitive functions by TGR63 in APP/PS1 AD phenotypic mice. (**A**) Tracing of vehicle and TGR63 treated mice (WT and AD) locomotion during open field (OF) test (test period: 5 min). (**B**) Total distance traveled by experimental mice cohorts. (**C**) Average number of entries into the center zone. (**D**) Distance traveled by experimental mice cohorts in the center zone. Data are presented as mean ± SEM, WT vehicle group N= 8, WT TGR63 group N= 8, AD vehicle group N= 8 and AD TGR63 group N= 9. (**E**) The novel object identification (NOI) test protocol: Image of experimental arenas during habituation, familiarization and test days. (**F** and **G**) The recognition of novel objects compared to old object on test day 1 and 2, respectively. Data are presented as discrimination index (DI) [DI= (time exploring the novel object – time exploring the familiar) / (time exploring novel + familiar) * 100], WT vehicle group N= 8, WT TGR63 group N= 8, AD vehicle group N= 8 and AD TGR63 group N= 8. (**H**) The Morris water maze (MWM) test analysis: Trajectory of experimental (vehicle and TGR63 treated) WT and AD mice in training period (day 1 and 4). (**I**) Latency time (second) of each cohort for searching the hidden platform during training. (**J**) The representative trace of experimental mouse in probe trail (no platform). (**K**) Percentage of total exploration by each cohort in target quadrant (platform was placed during training) and other quadrants in probe trial. (**L**) Average number of target (platform) crossing by each cohort during probe trail (no platform). Data are presented as mean ± SEM, WT vehicle group N= 10, WT TGR63 group N= 10, AD vehicle group N= 8 and AD TGR63 group N= 10. * p < 0.05, analyzed by two-way ANOVA followed by Bonferroni test.

The spatial and episodic memory formation under AD conditions were investigated through spatial learning and memory development tasks in MWM test (*34*). MWM test was performed in a water pool (radius: 70 cm) and experimental mice were trained four times in a day to find a hidden platform, which was removed in probe trial to assess the spatial memory (Fig. S15-18). The latency time to reach the hidden platform during the training period was recorded to determine spatial learning (Fig. 5H). As anticipated, AD vehicle cohort required more time (~ 70, 60 and 43 s) to reach the platform during training days (2^nd^, 3^rd^ and 4^th^, respectively), while other cohorts showed a smooth spatial memory formation with time (Fig. 5I). AD TGR63 cohort behaved like a healthy WT mouse and exhibited significant improvement in spatial memory formation compared to AD vehicle cohort. In the probe trial, AD vehicle cohort spent most of the time (~87% of total time) in other quadrants (without platform), while other cohorts (WT vehicle, WT TGR63 and AD TGR63) spent only ~67%, 58% and 66% of total time in without platform quadrants, respectively. The AD vehicle cohort spent minimum time (~13% of total time) in target quadrant (with platform) compared to WT vehicle cohort (~33% of total time). TGR63 does not affect the spatial memory formation and retrieval in the healthy brain, as the WT TGR63 cohort showed similar exploration (<35% of total time) tendency like WT vehicle cohort. Interestingly, AD TGR63 cohort explored <20% (~34% of total time) in the target quadrant than AD vehicle cohort, which is similar to that of healthy mice (Fig. 5K). Further, we determined the spatio-temporal memory by analyzing their activity in the platform region, which revealed AD vehicle cohort crossed the platform for minimum times (~1 time) compared to the WT vehicle cohort (~4 times) (Fig. 5L). Remarkably, AD TGR63 cohort crossed the platform region ~4 times, which is greater than the AD vehicle cohort. MWM study demonstrated significant effect of TGR63 treatment on the medial entorhinal cortex and hippocampus in the AD brain, the key areas for the development of spatial learning and memory. Overall, the significant enhancement of memory and cognitive performance in the behavioral studies is in excellent agreement with the amelioration of amyloid burden and associated neuronal toxicity in the AD (APP/PS1) mice validated the anti-AD credentials of TGR63.

## DISCUSSION

Several therapeutic candidates have been developed to modulate AD progression with Aβ burden as a therapeutic target (*5, 8, 35–37*). Identification of potent small molecules to ameliorate Aβ burden and associated cognitive deficits eluded researchers and clinicians to find an effective treatment for AD (*8*). Targeting Aβ burden includes the modulating production, misfolding and parenchymal plaques deposition of Aβ, promotion of plaques clearance and amelioration of neuropathological hallmarks and cognitive decline (*9, 18, 37*). Inhibition of β or γ-secretases is a promising approach to reduce the Aβ production, albeit their expression and functional relevance in others part of the body failing a large number of clinical candidates due to severe side effects and off-target interactions (*38*). The immunotherapy or acceleration of Aβ clearance strategies shown significant enhancement in the brain inflammatory response besides reduction in the cerebral amyloid burden and associated cognitive impairment. These observations and findings have reiterated the fact that targeting parenchymal plaques deposition and associated neurotoxicity through the meticulous design of small molecule inhibitors is a promising approach to develop a potential therapeutic candidate for the treatment of AD (*2, 8, 9, 39*). We designed a focused set of small molecules to identify a lead candidate to ameliorate Aβ burden and related neuropathological hallmarks to improve cognitive functions in AD mice model.

The sequential proteolytic cleavages of APP by β and γ-secretases produce Aβ peptides of variable lengths (36-43 amino acids) (*2, 40*). Among these Aβ peptides, Aβ42 is highly aggregation-prone and undergoes misfolding and ordered the assembly to form neurotoxic amyloid plaques which contribute to multifaceted toxicity including plasma membrane disruption, synaptic dysfunction, memory impairment, cognitive decline and neuronal loss in the AD brain (*47–47*). The modulation of severe amyloid burden and associated neurotoxicity to improve cognitive functions is a gigantic challenge to research and clinical community. There is an unmet need to develop a new class of efficient modulators of Aβ aggregation and related neurotoxicity through unique and robust drug design strategy (*9*). Here, we discuss a simple yet eloquent design of focused set of NMI-based small molecules and their structure-function relationship study to identify a lead candidate (TGR63) as *in vitro* and *in vivo* modulator of Aβ aggregation to tackle amyloid burden associated neuropathological hallmarks to ameliorate cognitive deterioration.

NMI-based small molecules (TGR60-65) were designed through systematic variation of substituents to fine tune the hydrophobicity and hydrophilicity balance required to interact and effectively modulate Aβ aggregation. A detailed *in vitro* biophysical and screening assays revealed NMI-core with of *N,N*-dimethylamine and *N,N,N*-trimethylpropan-1-aminium substituents (TGR63) emerged as an efficient inhibitor of Aβ aggregation. TGR63 was obtained by functionalizing 4-bromo-NMA with *N,N*-dimethylamine using Sonogashira coupling protocols followed by the conjugation of *N,N,N*-trimethylpropan-1-aminium as imide substituent (Scheme S1). The detailed evaluation by ThT fluorescence, dot blot, AFM and TEM analysis validated effective *in vitro* modulation of Aβ42 aggregation by TGR63 (Fig. 1 and 2). NMR study revealed molecular level interactions of TGR63 with Aβ42. A clear splitting pattern and downfield shift of aromatic protons (6.45–6.65 and 6.85–7.15 to 6.50–6.70 and 6.90–7.20, respectively) in ^1^H NMR spectra in the presence of Aβ42 revealed molecular interactions of TGR63 with Aβ42, which provided as a possible molecular mechanism of aggregation modulation (Fig. 1D). In addition, *in silico* results are in good agreements with experimental results and established the superiority of TGR63 in Aβ aggregation modulation. TGR63 efficiently binds with existing sites (core and surface binding) and an additional cryptic site (core binding site) of the amyloid fibril. It generates stable TGR63-Aβ complex through the strong van der Waals and electrostatic interactions. Interestingly, this stable complex formation significantly decreases the crucial interactions (salt bridge and hydrogen bonding) within amyloid fibrils and is proposed as a plausible mechanism behind its effective aggregation modulation. A recent study revealed that Aβ aggregation process promotes misfolded Aβ-membrane interactions and internalization of Aβ, which initiates various cell signaling cascades and interrupt physiological neuronal functions (*8*). We have shown the interaction of Aβ aggregation species on the plasma membrane and associated neuronal loss in cultured cells (*29*). Interestingly, modulation (inhibition and dissolution) of Aβ42 aggregation in presence of TGR63 reduced the membrane toxicity and rescued cultured neuronal cells.

As discussed (vide supra), the chronic Aβ plaques deposition induced dendritic and axonal atrophy in the AD brain contributing to the loss of mature neurons and neuronal circuit disruption (*10, 11, 48*). The soluble Aβ aggregation species interact with synaptic receptors (NMDA and AMPA) at the synaptic cleft hampering the neuronal signaling cascade, memory formation and cognitive functions (*12, 14, 49*). Double transgenic APP/PS1 AD mice show AD phenotypes viz., accumulation of chronic Aβ plaques, memory impairments, cognitive decline and neuronal loss with age (*4, 50*). Human transgenes APP and PS1 with Swedish and L166P mutations, respectively, are overexpressed in the APP/PS1 mouse brain and promote amyloidogenic APP cleavage to generate excess Aβ in CNS (*2, 30*). Accumulation of Aβ plaques has been supported by the postmortem report of the AD brains, and our immunohistochemistry data of the APP/PS1 mouse brain tissue fully supported Aβ plaque deposits in abundance. Deposition of Aβ plaques in the brain is the characteristic neuropathological hallmark of AD which subsequently cause neuropsychiatric dysfunction, as validated by the behavioral (OF) assay (Fig. 5). The downstream effects of Aβ burden associated cognitive dysfunction include interruption of neuronal circuits, synapse and synaptic plasticity, which result in deterioration of recognition ability, learning ability and spatiotemporal memory formation. Our NOR and MWM behavioral tests validated cognitive decline in APP/PS1 mouse and AD phenotype to evaluate the efficacy of our lead candidate.

The *in vivo* efficacy of TGR63 was evaluated in APP/PS1 mice model, and the results showed global improvement of cognitive and memory functions under progressive AD conditions. The trajectory of TGR63 treated AD mice in an unexplored arena (OF test) revealed improved locomotor and anxiety similar to that of vehicle-treated WT cohort, (Fig. 5A-D) confirming the therapeutic potential of TGR63 treatment. Acquisition, consolidation and retrieval of memory were assessed by the ability of TGR63 treated AD mice to discriminate the novel from the familiar object. TGR63 treated AD mice showed significantly improved discrimination of novel from the familiar object was similar to that of vehicle-treated WT cohort and the NOI (Fig. 5E-G). The NOI test further confirmed the effect of TGR63 in rescuing memory under progressive AD conditions. The learning ability and the acquisition of working and spatiotemporal memory of TGR63 treated AD mice was evaluated using standard MWM test. The MWM test results showed recovery of the learning, and spatiotemporal and working memory formation in TGR63 treated AD mice (Fig. 5HL). We hypothesize that the improved physiological brain functions of TGR63 treated AD mice are similar to healthy mice indicated rescue of synaptic plasticity upon TGR63 administration. The cognitive improvement was supported by the significant reduction in amyloid deposits in the APP/PS1 AD mice brain, as revealed by the postmortem immunohistochemical studies (Fig. 4). AD positive transgenic mice showed a significant deposition of Aβ plaques in different regions of the brain, including cortex and hippocampus. The observed amyloid deposits and associated cognitive decline directly correlates with AD phenotypes (*4*). The histochemical studies of TGR63 treated APP/PS1 AD mice showed a significant reduction of Aβ plaques throughout the brain, particularly hippocampal and cortical regions. The reduced neuropathological hallmark in the AD brain diminished the amyloid toxicity such as membrane disruption, synaptic dysfunction and neuronal loss, thereby improving the physiological brain functions (Fig. 5). Further, the pharmacokinetics study established serum stability, BBB permeability and biocompatibility of TGR63. The gold standard H&E staining of organs (heart, liver, kidney and spleen) from TGR63 treated AD and WT mice revealed biocompatibility that established TGR63 as a suitable candidate for prolonged administration through IP injection (Fig. 3).

In conclusion, the misfolding and aggregation of Aβ peptides into toxic soluble and insoluble aggregation species are hallmarks of AD progression and associated multifaceted toxicity. Accumulation of Aβ plaques in the brain directly correlates with AD phenotypes such as neuropsychiatric symptoms, learning deficiency, memory impairment, and cognitive decline. Modulation of Aβ burden and amelioration of associated neuropsychiatric symptoms are considered as the major therapeutic routes to treat AD. In this context, we designed, synthesized and identified a small molecule modulator of Aβ aggregation to ameliorate *in vitro* and *in vivo* Aβ induced neuronal toxicity and associated neuropsychiatric symptoms. The *in vitro* and *in cellulo* studies demonstrated that NMI derivative TGR63 with 4-ethynyl-*N,N*-dimethylaniline and *N,N,N*-trimethylethylenediamine functionalities bestowed right hydrophobicity-hydrophilicity balance to inhibit Aβ42 aggregation and associated neuronal toxicity. The detailed NMR and *in silico* study provided valuable insights on the molecular level interactions between TGR63 and Aβ species (monomers and fibrils), which revealed the plausible mechanism of aggregation inhibition and justified our design strategy. The *in vivo* pharmacokinetics study established serum stability, BBB permeability, *in cellulo* and *in vivo* biocompatibility and suitability of TGR63 for prolonged treatment through IP injection. TGR63 treated APP/PS1 mice brain tissue revealed a significant reduction of Aβ deposits validating its therapeutic efficacy as *in vivo* modulator of amyloid burden under progressive AD conditions. The treatment of APP/PS1 mice with TGR63 showed amelioration of learning deficiency, memory impairment, and cognitive decline as revealed by distinct OF, NOI and MWM behavioral tests. Remarkably, the improvement in brain functions (learning efficiency, memory formations and cognitive functions) under progressive disease conditions is in excellent correlation with the reduced cortical and hippocampal Aβ load following theTGR63 treatment. These key attributes have validated the potential of TGR63 as a promising candidate for the treatment of AD.

## MATERIALS AND METHODS

### Synthesis of TGR63

To a solution of 4-((4-*N,N* dimethylaniline) ethynyl)-1,8-naphthalic anhydride (200 mg, 0.58 mmol) dispersed in isopropanol, DIPEA (31 mL, 1.7 mmol) and 2-amino-*N,N,N*-trimethylethanaminium (60 mg, 0.58 mmol) were added and refluxed at 80 °C for 6 h. The reaction mixture was extracted with ethyl acetate, washed with brine, dried over Na_2_SO_4_ and evaporated to obtain the crude product. The crude product was purified by column chromatography on silica gel using 1% MeOH in CHCl_3_ as an eluent to afford a red coloured solid in good yield (75%). ^1^H NMR (DMSO *d*_6_, 400 MHz) δ 8.60 (d, 1H, *J* = 0.8), 8.58 (d, 1H, *J* = 1.2), 8.48 (d, 1H, *J* = 7.6), 8.03 (d, 2H, *J* = 4.4), 8.01 (t, 2H, *J* = 4.8), 7.61 (d, 2H, *J* = 2), 6.80 (d, 2H, *J* = 8.8), 4.48 (t, 2H, *J* = 13.6), 3.65 (t, 2H, *J* = 14.8), 3.21 (s, 9H), 3.01 (s, 6H); ^13^C NMR (DMSO *d*_6_, 100 MHz) δ 163.3, 163, 158, 150.9, 133.2, 132.5, 131.3, 130.6, 130.4, 129.7, 128, 127.6, 122.4, 120.5, 111.8, 106.9, 102.3, 85.9, 52.4, 33.6; HRMS (ESI-MS): found 426.2176, calcd. for C_27_H_28_N_3_O_2_ [M]^+^ m/z = 426.2176.

## SUPPLEMENTARY MATERIALS

Materials and Methods

Scheme S1. Syntheses schemes for TGR60-65

Fig. S1. Inhibition and dissolution of Aβ42 aggregates studied by ThT assay

Fig. S2 and S3. Neuronal rescue

Table S1-3. *In silico* analysis

Fig. S4. *In silico* analysis

Fig. S5. The calculation of lethal dose 50 (LD50) of TGR63 by intraperitoneal administration

Fig. S6 and S9. MALDI analysis

Fig. S7. *In vitro* serum stability of TGR63

Fig. S8. *In vitro* calculation of LogP

Fig. S10. Toxicology study of TGR63 in AD mice

Fig. S11-14. The locomotion of vehicle treated WT mice cohort during OF test.

Fig. S15-18. The trajectory of vehicle treated WT mice cohort during MWM probe trail (without platform)

Video 1. Representative video of OF test

Video 2. Representative video of NOI test

Video 3. Representative video of MWM test

Data file S1. Characterization data of TGR60-65

## Acknowledgments

We thank JNCASR, Prof. A. Anand, Dr. R. G. Prakash, and JNCASR animal facility. We also thank Dr. H. K. Gowda (Karnataka Veterinary, Animal and Fisheries University, Karnataka) for helping in toxicology study.

## Author contributions

K.R., S.S. and T.G. designed the project. K.R. and S.S. synthesized compounds and performed photophysical, biophysical and *in vitro* studies. S.S. M.R. and S.A. performed *in vivo* and behavioral studies, S.S. and D.S. preformed post-administration immunohistological experiments. S.S carried out the pharmacokinetics analysis. N.A.M performed all the computational studies involving molecular docking and molecular dynamics. The *in vivo* behavior data was analyzed and discussed with J.P.C. and T.G., T.G. supervised overall the project. S.S. and T.G. prepared the manuscript and all the authors read and commented.

## Data and materials availability

All data related to this study can be found in the paper or the Supplementary Materials.

