## Supplementary Materials for "Naphthalene monoimide derivative ameliorates amyloid burden and cognitive decline in a transgenic mouse model of Alzheimer’s disease"

**This PDF file includes:**

Scheme S1. Syntheses schemes for TGR60-65

Fig. S1. Inhibition and dissolution of Aβ42 aggregates studied by ThT assay

Fig. S2 and S3. Neuronal rescue

Table S1-3. In silico analysis

Fig. S4. In silico analysis

Fig. S5. The calculation of lethal dose 50 (LD50) of TGR63 by intraperitoneal administration

Fig. S6 and S9. MALDI analysis

Fig. S7. In vitro serum stability of TGR63

Fig. S8. In vitro calculation of LogP

Fig. S10. Toxicology study of TGR63 in AD mice

Fig. S11-14. The locomotion of vehicle treated WT mice cohort during OF test.

Fig. S15-18. The trajectory of vehicle treated WT mice cohort during MWM probe trail (without platform)

Video 1. Representative video of OF test

Video 2. Representative video of NOI test

Video 3. Representative video of MWM test

Data file S1. Characterization data of TGR60-65

References (1-9)

**General methods**.

All solvents and reagents were obtained from Spectrochem or Merck and used without any further purification unless mentioned. Dulbecco's Modified Eagle Medium/Nutrient Mixture F 12 (DMEM F12), Roswell Park Memorial Institute (RPMI), fetal bovine serum (FBS) and horse serum (HS) was obtained from Invitrogen. Argon or nitrogen atmosphere was maintained for all the reactions. Agilent Cary series UV−Vis-NIR absorption, Agilent Cary eclipse fluorescence spectrophotometers and microplate reader (SpectrsMax i3x) were used to perform absorption and fluorescence assay. $^{1}$H and $^{13}$C NMR spectra were recorded in Bruker AV−400 and JEOL-600 MHz spectrometers, and tetramethylsilane (TMS) was used as an internal standard. All the raw data was processed and analyzed using Prism 6 or Origin 8.5 software. HRMS spectra were acquired using Agilent 6538 UHD HRMS/Q-TOF high-resolution spectrometer. The calculated amount of inhibitors were dissolved in deionized water (Milli Q) (contain 5% dimethyl sulfoxide) to store (-20 °C) and diluted in phosphate buffered saline (pH= 7.4) for the experiments. Amyloid beta peptide was obtain from Merck (PP69-0.05 MG). The anti-amyloid fibrils (OC) and oligomers (A11) specific primary antibodies were obtained from Merck and ThermoFisher, respectively, to performed immunohistochemistry. Thioflavine T (ThT) was obtain from Sigma-Aldrich (T3516) and CQ probe was obtained from VNIR Biotech. Sodium citrate buffer was obtained from Fisher Scientific, India (6132-4-3,) and DAPI (4′,6-diamidino-2-phenylindole) was obtain from Vector Laboratories, CA, USA (H-1200). Blue Star micro slides were used to mount the brain sections. The tissue homogenizer (D9938) and primers were obtained from Sigma-Aldrich. We got the mice ear tagging set from Jaxson laboratory, USA. The experimental brains were sectioned using Leica Vibratome (VT1200). All the images (cells and the brain) were captured using confocal fluorescence microscope (Olympus FV3000).

**Synthesis of 4-((4-*N,N* dimethylaniline) ethynyl)-1,8-naphthalic anhydride**.

To a solution of 4-bromo-1,8-naphthalic anhydride (200 mg, 0.72 mmol) in dimethyl formamide (DMF)/triethylamine ($Et_3N$) (1 : 1) under argon atmosphere, $Pd(PPh_3)_4$ (27 mg, 23 µmol), sodium ascorbate (10 mg, 50 µmol), copper (II) sulfate (2 mg, 8 µmol) and 4-ethynylanisole (93.6 µL, 0.72 mmol) were added. The reaction mixture was stirred for 4 h at 80 °C and completion of the reaction was monitored by thin layer chromatography (TLC). The reaction mixture was extracted into ethyl acetate, washed with $NH_4Cl$ and brine, dried over $Na_2SO_4$ and evaporated under vacuo to obtain the crude product. The product was re-dissolved in ethyl acetate, precipitated with diethyl ether and the pure product (1) was collected by filtration. The product was obtained as dark red coloured solid in good yield (68%). $^1$H NMR ($CDCl_3$, 400 MHz) δ 8.65 (d, 2H, *J* = 6.4), 8.64 (d, 2H, *J* = 4.2), 8.55 (d, 2H, *J* = 8), 7.92 (d, 2H, *J* = 7.6), 7.89 (t, 2H, *J* = 15), 7.55 (d, 2H, *J* = 4.2), 6.73 (d, 2H, *J* = 6.4), 3.06 (s, 6H) ; $^{13}$C NMR ($CDCl_3$, 100 MHz) δ 163.8, 151, 134.2, 133.6, 132.7, 131.5, 130.7, 130.4, 130, 127.4, 116.5, 111.7, 108, 40.1; HRMS (ESI-MS): found 342.1145, calcd. for $C_{22}H_{16}NO_3$ $[M+H]^+$ m/z = 342.1112.

**Synthesis of TGR60**.

To a solution of naphthalic anhydride (114 mg, 0.58 mmol) dispersed in isopropanol, *N,N*-diisopropylethylamine (DIPEA; 31 µL, 1.7 mmol) and 2-amino-*N,N,N*-trimethylethanaminium (60 mg, 0.58 mmol) were added and refluxed at 80 °C for 6 h. The reaction mixture was extracted with chloroform ($CHCl_3$), washed with brine, dried over $Na_2SO_4$ and evaporated under vacuo to obtain the crude product. The crude product was purified by column chromatography on silica gel using 2% methanol (MeOH) in $CHCl_3$ as an eluent to afford a white solid in excellent yield (88%). $^1$H NMR (DMSO $d_6$, 400 MHz) δ 8.54-8.50 (m, 4H), 7.93-7.89 (m, 2H), 4.49 (t, 2H, *J* = 14.4), 3.66 (t, 2H, *J* = 14.4), 3.23 (s, 9H); $^{13}$C NMR (DMSO $d_6$, 100 MHz) δ 163.4, 134.7, 131.3, 130.9, 127.4,

127.3, 121, 89, 61.9, 52.5, 33.6; HRMS (ESI-MS): found 283.1439, calcd. for $C_{17}H_{19}N_2O_2$ $[M]^+$ m/z = 283.1441.

**Synthesis of TGR61**.

To a solution of 4-dimethylamine-1,8-naphthalic anhydride (139 mg, 0.58 mmol) dispersed in isopropanol, DIPEA (31 µL, 1.7 mmol) and 2-amino-*N*,*N* -trimethylethanaminium (60 mg, 0.58 mmol) were added and refluxed at 80 °C for 6 h. The reaction mixture was extracted with $CHCl_3$, washed with brine, dried over $Na_2SO_4$ and organic layer was evaporated to obtain the crude product. The crude product was purified by column chromatography on silica gel using 3.5% MeOH in $CHCl_3$ as an eluent to afford a yellow solid in good yield (54%). $^1H$ NMR (DMSO $d_6$, 400 MHz) δ 8.50 (d, 1H, $J$ = 6.4), 8.49 (d, 1H, $J$ = 4.2), 8.38 (d, 1H, $J$ = 8.4), 7.80 (d, 1H, $J$ = 7.2), 7.78 (d, 1H, $J$ = 7.2), 7.24 (d, 1H, $J$ = 4.2), 4.96 (t, 2H, $J$ = 14), 3.64 (t, 2H, $J$ = 14), 3.20 (s, 9H), 3.12 (s, 6H); $^{13}C$ NMR (DMSO $d_6$, 100 MHz) 163.7, 162.9, 156.9, 132.6, 132.1, 130.8, 124.9, 124, 122, 112.8, 112.5, 52.4, 44.3, 33.4; HRMS (ESI-MS): found 326.1864, calcd. for $C_{19}H_{24}N_3O_2$ $[M]^+$ m/z = 326.1863.

**Synthesis of TGR62**.

To a solution of 4-(benzylethynyl)-1,8-naphthalic anhydride (172 mg, 0.58 mmol) dispersed in isopropanol, DIPEA (31 µL, 1.7 mmol) and 2-amino-*N,N,N*-trimethylethanaminium (60 mg, 0.58 mmol) were added and refluxed at 80 °C for 6 h. The reaction mixture was extracted with $CHCl_3$, washed with brine, dried over $Na_2SO_4$ and evaporated under vacuo to obtain the crude product. The crude product was purified by column chromatography on silica gel using in $CHCl_3$ as an eluent to afford a yellow solid in good yield (74%). $^1H$ NMR (DMSO $d_6$, 400 MHz) δ 8.83 (d, 1H, $J$ = 8.4), 8.61 (d, 1H, $J$ = 7.2), 8.51 (d, 1H, $J$ = 7.6), 8.13 (d, 1H, $J$ = 7.6), 8.05 (t, 1H, $J$ = 15.6), 7.81-7.78 (m, 2H), 7.54-7.52 (m, 3H), 4.49 (t, 2H, $J$ = 14.4), 3.67 (t, 2H, $J$ = 14.4), 3.24 (s, 9H); $^{13}C$

NMR (DMSO $d_6$, 100 MHz) 163.2, 162.9, 132.3, 131.9, 131.4, 131, 130.9, 130.2, 129.9, 128.9, 128.4, 127.4,5, 126.5, 122.5, 121.8, 121.2, 99, 86.0, 61.9, 54.8, 52.4, 33.7; HRMS (ESI-MS): found 384.1693, calcd. for $C_{25}H_{23}N_2O_2$ $[M]^+$ m/z = 384.1854.

**Synthesis of TGR64**.

To a solution of 4-((4-*N*,*N* dimethylaniline) ethynyl)-1,8-naphthalic anhydride (200 mg, 0.58 mmol) dispersed in isopropanol, DIPEA (31 μL, 1.7 mmol) and tert-butyl 2-aminoethylcarbamate (39 mg, 0.58 mmol) were added and refluxed at 80 °C for 6 h. The reaction mixture was extracted with ethyl acetate, washed with brine, and dried over $Na_2SO_4$. The crude product was purified using column chromatography on silica gel using 0.25% MeOH in $CHCl_3$ as an eluent to afford a red coloured solid. Then the compound was deprotected using TFA (95% TFA, 4.5% DCM and 0.5% TIPS) and the product was precipitated to obtain pure product in good yield (68%). $^1H$ NMR (DMSO $d_6$, 400 MHz) δ 8.77 (d, 1H, *J* = 1.8), 8.56 (d, 1H, *J* = 3.6), 8.45 (d, 1H, *J* = 3.8), 8.00 (d, 2H, *J* = 3), 7.97 (d, 2H, *J* = 8.8), 7.59 (d, 2H, *J* = 3.3), 6.74 (d, 2H, *J* = 3.8), 4.33 (t, 2H, *J* = 11.6), 3.17 (s, 2H), 3.01 (s, 6H); $^{13}C$ NMR (DMSO $d_6$, 100 MHz) δ 163.8, 163.5, 150.8, 133.2, 132.2, 131.1, 130.5, 130.1, 129.7, 127.9, 127.7, 122.7, 120.8, 111.8, 107, 102, 85, 37.6, 37.5; HRMS (ESI-MS): found 383.1767, calcd. for $C_{24}H_{21}N_3O_2$ $[M]^+$ m/z = 383.1634.

**Synthesis of TGR65**.

To a solution of 4-((4-*N*,*N* dimethylaniline) ethynyl)-1,8-naphthalic anhydride (200 mg, 0.58 mmol) dispersed in isopropanol, DIPEA (31 μL, 1.7 mmol) and 2-(2-aminoethoxy)ethanol (22 mL, 0.58 mmol) were added and refluxed at 80 °C for 6 h. The reaction mixture was extracted with ethyl acetate, washed with brine, and dried over $Na_2SO_4$ and evaporated under vacuo to obtain the crude product. The crude product was purified by column chromatography on silica gel $CHCl_3$ as an eluent to afford a red coloured solid in good yield (72%). $^1H$ NMR (DMSO $d_6$, 400 MHz) δ 8.76

(d, 1H, $J$ = 8.4), 8.55 (d, 1H, $J$ = 7.2), 8.43 (d, 1H, $J$ = 7.6), 7.98 (d, 2H, $J$ = 8), 7.95 (t, 2H, $J$ = 1.6), 7.59 (d, 2H, $J$ = 8.8), 6.79 (d, 2H, $J$ = 8.8), 4.25 (t, 2H, $J$ = 12.8), 3.67 (t, 2H, $J$ = 12.8), 3.47 (s, 4H), 3.31 (s, 4H), 3.00 (s, 6H); $^{13}C$ NMR (DMSO $d_6$, 100 MHz) 163.2, 162.9, 133.2, 132.1, 131.1, 130.5, 130.2, 129.7, 127.9, 127.6, 127.5, 122.4, 120.6, 111.8, 107, 101.8, 85, 72, 66.8, 60.1, 28.9; HRMS (ESI-MS): found 429.1803, calcd. for $C_{26}H_{25}N_2O_4$ $[M+H]^+$ m/z = 429.1814.

**Preparation of Aβ42 aggregation species.**

Aβ42 peptide was dissolved in hexafluoro-2-propanol (HFIP, 250 μL) and incubated for 1 h at room temperature, and HFIP was removed by nitrogen gas flow. The processed Aβ42 peptide was dissolved in 2% DMSO or NaOH solution (100 mM) containing PBS buffer (pH = 7.4) to prepare the monomaric Aβ42 solution and the peptide concentration was calculated by UV-visible absorbance study ( $\varepsilon = 1450\ cm^{-1}\ M^{-}1$). Oligomers were prepared by incubating Aβ42 monomers for 24 h at 4 ºC. Similarly, Aβ42 monomers were incubated for 2 days in PBS buffer (pH = 7.4) at 37 ºC to prepare fully grown fibrillar aggregates and the presence of Aβ42 fibrils was confirmed by ThT assay.

**Dot blot analysis.**

To demonstrate the Aβ42 aggregation modulation ability of TGR63, we performed dot blot analysis. The freshly prepared Aβ42 (10 μM) sample was incubated (at 4 and 37 ºC) with TGR63 and alone independently for 24 h (oligomers) and 48 h (fibrils) without shaking, respectively. The incubated samples were dotted on the PVDF membrane and allowed to dry. The PVDF membranes was blocked using 5% skimmed milk (HIMEDIA, GRM 1254) in PBS for 1 h at room temperature. The blots were washed (3 times) with 1% of Tween 20 (HIMEDIA GRM156) containing PBS

(PBST) for 10 min and incubated with A11 (1:1000) and OC (1:1000) primary antibody, specific to Aβ42 oligomers and fibrils, respectively, at 4 °C for 16 h. Then the unbound primary antibody was removed by PBST wash (3 times) and incubated with HRP conjugated anti-mouse (for A11) anti-rabbit (for OC) secondary antibody (Biorad, 1706515), which was diluted 10000 times. Further, nonspecific binding was removed with PBST wash and the blots were developed with the treatment of enhanced chemiluminescence (ECL) reagent in Versa Doc (Biorad) instrument.

**Cell culture.**

SHSY5Y and PC12 cells were cultured using DMEM/F-12 (Dulbecco's Modified Eagle Medium/Nutrient Mixture F 12) medium (Gibco, Invitrogen) containing 10% of FBS (fetal bovine serum) and 1% PS (pen-strep) and RPMI (Roswell Park Memorial Institute) medium (Gibco, Invitrogen) with fetal bovine serum (FBS, 10%), horse serum (HS, 5%), and pen-strep (1%), respectively, under the cell growing condition (37 °C temperature and 5% $CO_2$ atmosphere).

**Imaging of Aβ42 fibrils in cellular milieu.**

We performed Aβ42 fibrils imaging in SHSY5Y cells to study the plasma membrane toxicity of Aβ42 fibrils in absence and presence of TGR63.(*1*) For imaging experiment, cells were cultured in petri dishes (35 mm) and treated with TGR63 treated and untreated Aβ42 fibrils for 2 h. The experimental cells were washed and fixed with PBS and 4% PFA, respectively. Then, the cells were treated with red fluorescent-labeled ($\lambda_{ex}$= 633 nm and $\lambda_{em}$= 650 nm) secondary antibody or CQ followed by Aβ42 fibrils specific primary antibody, OC (1:250) and DAPI to capture images under the confocal fluorescence microscope.

**Neuronal cell rescue.**

To demonstrate the neuronal cells rescue ability of inhibitors (TGR60-65) form Aβ42 peptide toxicity, we performed MTT assay.(*2*) The cells were (15,000 cells/well) cultured in a 96-well plate using cell growing media and incubated with for Aβ42 peptide and inhibitors for 24 h at 37 °C temperature within 5% $CO_2$ atmosphere. Further MTT (5 mg/mL) solution was added into the experimental cell media and incubated for 2.5 h. Finally, the experimental medium was removed and 100 μL of DMSO:MeOH (1:1) mixture was added, and the absorbance (570 nm) was monitored using microplate reader.

***In silico* assessment.**

The molecular level interaction of compounds TGR60-65 with Aβ monomer and fibrils was studied using an integrated approach involving molecular docking, molecular dynamics and binding free energy calculations. The molecular docking approach was employed to identify all the low energy binding sites and modes for TGR63 with monomeric and fibril forms of Aβ. The target structure for Aβ monomer is based on the NMR structure deposited in the protein databank (pdb id is 1IYT) (*3*). The Aβ monomer structure reported corresponds to aqueous solvent environment and has two helical regions (involving residues in the range 8-25 and 28-38) with a type I β-turn. There are10 different models reported for Aβ monomer and the docking was carried out for all the models using auto dock software (*4*). Since the binding modes are not known, a blind docking procedure was adopted by incorporating the entire peptide within a sufficiently larger grid box having grid points 110, 90, 160 along x, y and z directions. The grid point spacing was chosen to the default value of 0.375 Å. The Lamarckian genetic algorithm was adopted to find out the least energy binding sites and binding modes for TGP63 within the Aβ peptide. The model with the least energy has been adopted for the subsequent molecular dynamics (MD) simulations. At least three independent least energy binding sites were observed. A complex structure of Aβ monomer with three molecules of TGR63 bound in the lowest energy binding modes was prepared as the input structure for MD. The simulation box was solvated with approximately 13350 water molecules. The MD simulations followed a routine protocol and involved minimization, simulation in constant volume ensemble and simulation in isothermal isobaric ensemble. A short time scale equilibration run was carried out to allow the system to evolve to the ambient temperature and pressure. Followed by the equilibration run for a time scale of 5 ns, final production run was carried out for a time scale of 40 ns. All MD simulations were performed using Amber16 software (*5*). The time step for solving the Newton's equation of motion was set to 2 fs. 500 configurations from last 5 ns were used for carrying out ensemble average of binding free energies using molecular mechanics-generalized Born surface area (MM-GBSA) approach. This approach involves calculations of free energies of complex and subsystems namely ligands and target. The binding free energies were computed as the difference between the complex and subsystem free energies. The solvents and ions were stripped from the simulation trajectories and only the receptor-ligand coordinates were used for the free energy calculations. To account for the solvent effect, the approach employs generalized Born approach and polar and non-polar solvation energies together account for the free energy changes associated with binding due to aqueous environment. A similar approach described above for Aβ monomer was adopted for Aβ fibril. The Aβ fibril structure reported in protein databank (pdb id is 5OQV) based on cryo-EM was used for molecular docking (*6*). This structure constitutes of 9 Aβ peptides organized with LS topology as shown in Figure2a and the c-terminal regions are protected from exposure to solvents. Grid box chosen for docking includes entire fibril structure such that all possible surface sites can be identified. The number of grid points along x, y and z directions were chosen as 220, 200 and 130 with a default grid spacing. The molecular docking study showed that TGR63 can bind to 6 different binding sites in Aβ fibril. The binding modes in the initial configuration for TGR63 are shown in Figure 1E and 1F. The binding modes for TGR64 and TGR65 are shown in Figure S3. The MD simulation for complex of Aβ fibril with TGR63 in different binding sites was carried out following the procedure described for Aβ monomer. The total time for the production run was 40 ns. The binding free energies for TGR63 in different binding sites of Aβ fibril were computed using MM-GBSA approach. The binding free energies for other molecules TGR60, TGR61, TGR62, TGR64, TGR65 were also computed following the protocols described above.

**Animal maintenance.**

Double transgenic mice [B6.Cg-Tg(*APP*$^{swe}$*,PSEN1dE9*)85Dbo/Mmjax] were obtained from the Jackson Laboratory (MMRRC stock no 34832) and maintained in JNCASR Animal Facility under 12 h light and dark cycle and were bred with WT mice (C57/BL6). All the animal maintenances and studies were performed according to the protocols of the Institutional Animal Ethics Committee (IAEC), JNCASR. The protocol (TG001) was approved by the IAEC and Committee for purpose of Control and Supervision of Experiments on Animals (CPCSEA), New Delhi.

**Genotyping of mice.**

All the mice were genotyped at 4-6 weeks of age. Genomic DNA was collected from the mouse tail and processed with NaOH and Tris-HCl buffer. The Jackson Laboratory's protocol was followed to confirm the Alzheimer's positive mice (*7*).

The following primer sequences used for genotyping:

APP:5’-GACTGACCACTCAGCCAGGTTCTG-3’;

5’CTTGTAAGTTGGATTCTCATATCCG-3’;

PSEN1: 5’-ATTAGAGAACGGCAGGAGCA-3’; 5’-GCCATGAGGGCACTAATCAT-3’.

**Blood brain barrier (BBB) crossing and serum stability experiment.**

Vehicle (PBS) and TGR63 were administrated to WT mice and sacrificed at different time points to collect the blood and brain samples. Collected blood and brain samples were processed to obtain blood serum and brain lysate, which were analyzed through MALDI mass using α-cyano-4-hydroxycinnamic acid (CCA) matrix for identifying TGR63 in blood serum and brain.

**Immunohistochemistry.**

After all the behavioral studies, experimental AD and WT mice were sacrificed (cervical dislocation) and their brains were dissected out. The collected brains were fixed with paraformaldehyde (4%) for 48 h and then rehydrated with 30% sucrose solution. The experimental brains were sectioned (40 μm) using vibratome (Leica), and stored at -20 °C. The brain sections were treated with 10 mM sodium citrate buffer to remove the antigen. The sections were blocked with Bovine Serum Albumin (BSA, 2%) and goat serum (1%) for 4 h followed by permeabilization with 0.1% Triton-X-100 containing 0.1 M PBS (PBTx). After blocking, the sections were incubated with primary antibodies for 24 h at 4 °C on a shaker (slow speed). The unbound primary antibody was washed out and the sections were incubated with fluorescently (green and red) labeled secondary antibodies for 4 h at room temperature in the dark on a shaker. Finally, these brain sections were incubated with DAPI for 10 min and taken on glass slide for mounting with vecta-sheild and imaged using confocal fluorescence microscope. ImageJ was used to analyze the images and for quantification of results.

**Open-Field test.**

This experiment was conducted in a 45 × 45 x 45 cm OF apparatus made of grey plywood. All the mice were placed in a same position (corner) of the open field and the movement of the mouse was recorded for 5 min using a video camera fix on the top of the apparatus and analyzed using smart 3.0 software. 70% ethanol was used to clean the test apparatus after each and every experiment and left for 5 min for air dry.

**NOI test.**

The improvement of recognizing memory by TGR63 treatment were assessed using the NOI test as previously described (*8*). The experiment was performed using a 33 × 20 X 33 cm (LxHxW) OF platform. All the experimental mice were habituated for 1 day in the platform and two diagonally located identical objects were familiarized in the 1$^{st}$ day. During testing (after 24 and 48 h) one novel object was introduced in place of one familiar object, respectively and the mice were allowed to explore for 10 min. The exploration time of familiar and novel objects was recorded using stopwatch. The discrimination index (DI) [DI= (time exploring the novel object – time exploring the familiar) / (time exploring novel + familiar) * 100] was calculated to analyze the exploration tendency. After each and every experiment the test apparatus and objects were cleaned with 70% ethanol and wiped out with tissue paper.

**MWM test.**

Morris water maze experiment was performed with our experimental mice, which is extensively used to analyze the spatial memory and learning as previously described (*9*). This experiment was conducted in a tub of 122 cm in diameter and 90 cm in depth and the pool was filled with water for up to 60 cm. Non-toxic paint was used to make the water opaque and pool temperature was maintained at 26 ± 0.5 °C. A 14 $cm^2$ platform was placed in the center of South-West (SW) quadrant of the pool, which was submerged 1 cm from the surface. During all training sessions, the platform remained in the same position and removed from the pool in the probe test. Mice were placed into the pool facing the wall and allowed 60 s to find the hidden platform. If the mouse failed to find the hidden platform in given time, it was guided towards the platform and allowed to stay on it for 60 s before returning to its home cage. This procedure was repeated four times in a day, and the starting position was different each time. After every trial, the mouse was dried off with tissue paper and clean towel. On $5^{th}$ day, the probe test was performed by removing the hidden platform. Mice were placed on the opposite quadrant (NE) where the platform used to be (SW). All tested mice were allowed 60 s to find the platform in the entire pool. Videos were recorded using Nikon coolPex camera and analyzed using smart 3.0 software.

**Statistical analysis.**

All the results were plotted, and statics were performed using GraphPad Prism 6. Significance level between different groups were assessed using Two-way ANOVA with more than one independent variable. Further, significant difference was determined using Bonferroni's multiple comparisons Post hoc test ($^* p < 0.05$).

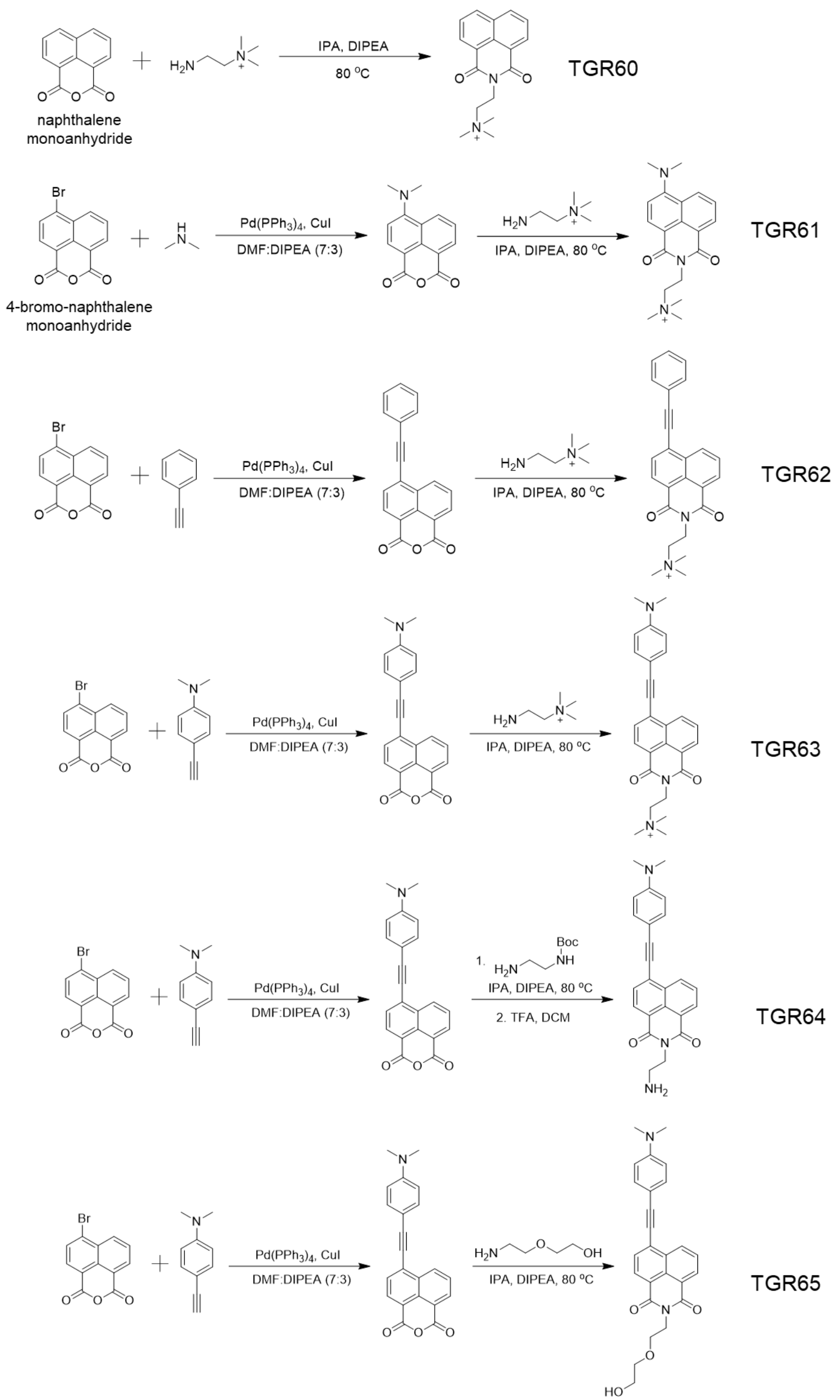


**Scheme S1.** Syntheses of small molecules TGR60-65 with NMI scaffold.

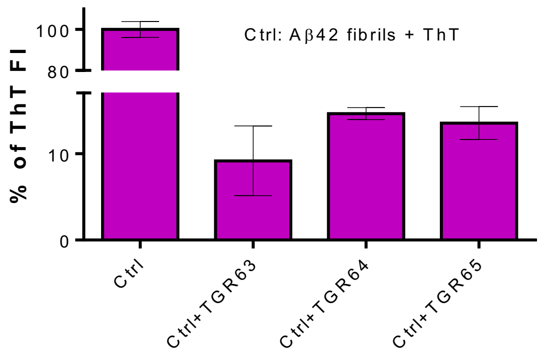


**Fig. S1.** Dissolution of Aβ42 aggregates studied by ThT assay. The data show the percentage (%) of ThT fluorescent intensity (FI) at 482 nm in presence of Aβ42 fibrils (10 μM) alone (Ctrl) and with inhibitors (30 μM). Each experiment was repeated three times (n = 3). Error bars represent the average±SEM of the fluorescence measurement.

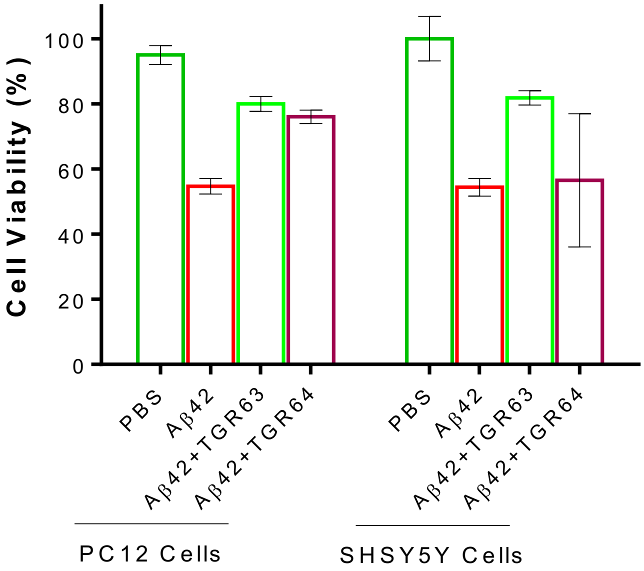


**Fig. S2.** *In vitro* neuronal rescue from Aβ42 toxicity by TGR63 and TGR64. The observed cell viability of cultured neuronal cells (PC12 and SHSY5Y) after independently incubating (24 h) with Aβ42 (20 μM) peptides in absence and presence of inhibitors (TGR63 and TGR64) in the 1:2 molar stoichiometric ratio.

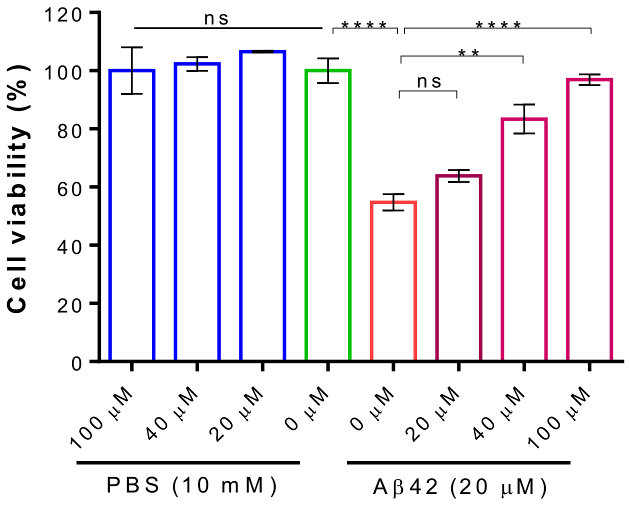


**Fig. S3.** *In vitro* neuronal rescue from Aβ42 toxicity by TGR63. The observed cell viability of cultured neuronal cells (SHSY5Y) after incubating (24 h) with different concentrations of TGR63 (20, 40 and 100 µM) in absence and presence of and Aβ42 (20 µM) peptides.

**Table S1.** Number of salt bridges and hydrogen bonds present in Aβ42 fibrils in the absence and presence of inhibitors (TGR63 and TGR64).

| System | Number of salt bridges | Number of hydrogen bonds |
|---|---|---|
| Aβ fibril in water | 48 | 81 |
| Aβ fibril + TGR63 in water | 41 | 75 |
| Aβ fibril + TGR64 in water | 54 | 74 |

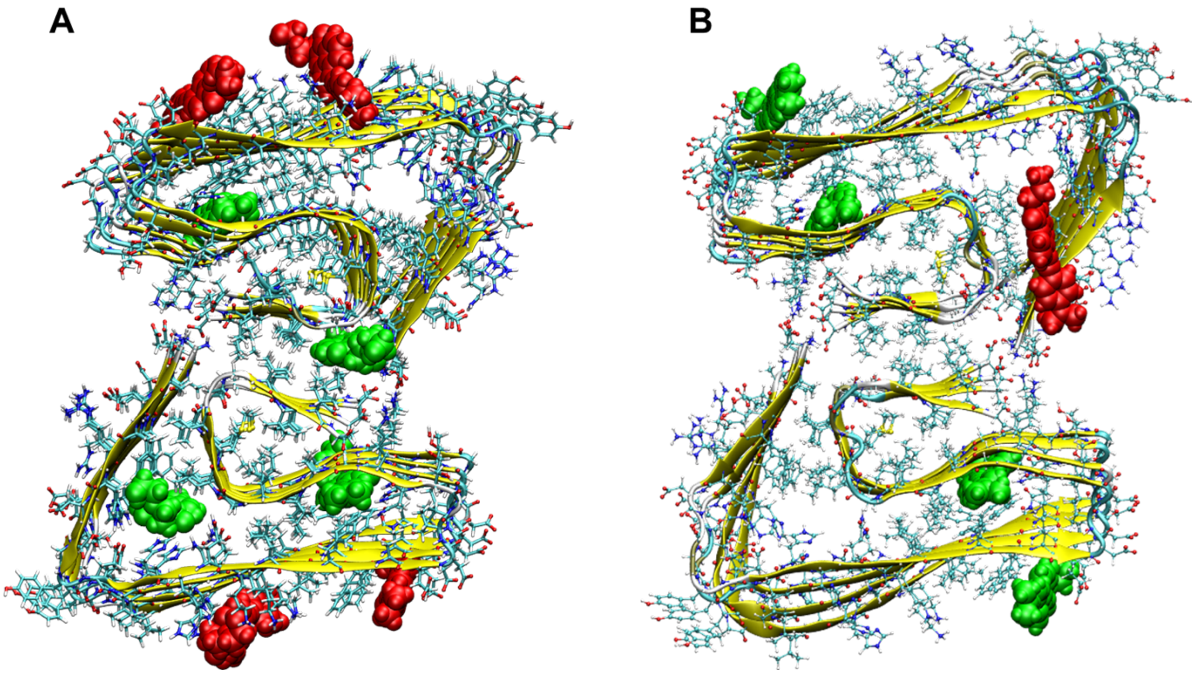


**Fig. S4.** Different binding sites for TGR64 (A) and TGR65 (B) in Aβ42 fibril.

**Table S2.** Binding free energies for TGR63 in different binding sites of Aβ42 fibril. Different contributions to binding free energies are provided. The energies are in kcal/mol and the standard errors in binding free energies are in the range 0.12 to 0.22 kcal/mol.

| Sites | $\Delta E_{vdw}$ | $\Delta E_{elec}$ | $\Delta E_{polar\ solvation}$ | $\Delta E_{non\text{-}polar\ solvation}$ | $\Delta G_{binding}$ |
|---|---|---|---|---|---|
| Site-1 | -71.8 | -310.1 | 338.1 | -6.8 | -50.6 |
| Site-2 | -74.4 | -309.1 | 342.2 | -6.5 | -47.8 |
| Site-3 | -41.0 | -377.8 | 388.3 | -4.8 | -35.4 |
| Site-4 | -36.2 | -361.5 | 371.8 | -4.4 | -30.3 |
| Site-5 | -38.2 | -203.5 | 221.4 | -4.1 | -24.4 |
| Site-6 | -23.4 | -317.8 | 329.8 | -3.1 | -14.5 |
| Site-7 | -51.1 | -301.3 | 323.4 | -6.0 | -34.9 |

**Table S3.** Binding free energies for TGR63 in different binding sites of Aβ42 monomer. The energies are in kcal/mol and the standard errors are in the range 0.1 to 0.3 kcal/mol.

| Sites | $\Delta E_{vdw}$ | $\Delta E_{elec}$ | $\Delta E_{polar\ solvation}$ | $\Delta E_{non\text{-}polar\ solvation}$ | $\Delta G_{binding}$ |
|---|---|---|---|---|---|
| Site-1m | -33.2 | -75.9 | 88.9 | -3.7 | -24.1 |
| Site-2m | -20.1 | -62.0 | 75.2 | -2.5 | -9.4 |
| Site-3m | -34.6 | -76.3 | 88.3 | -3.9 | -26.5 |

**A**

| TGR63 (mg/kg) | Number of mice | | | % of deaths |
|---|---|---|---|---|
| | Total | Lived | Died | |
| 179.0 | 5 | 2 | 3 | 60.00% |
| 56.0 | 5 | 5 | 0 | 0.00% |
| 17.50 | 5 | 5 | 0 | 0.00% |
| 5.5 | 5 | 5 | 0 | 0.00% |
| 1.7 | 5 | 5 | 0 | 0.00% |

**B**

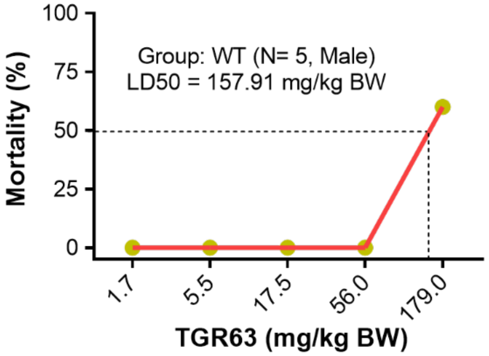


**Fig. S5.** The calculation of lethal dose 50% (LD50) of TGR63 through intraperitoneal administration. (**A**) Table of experimental details and the final observation on 14th day. (**B**) The mortality (%) is plotted against TGR63 concentration and calculation of LD50.

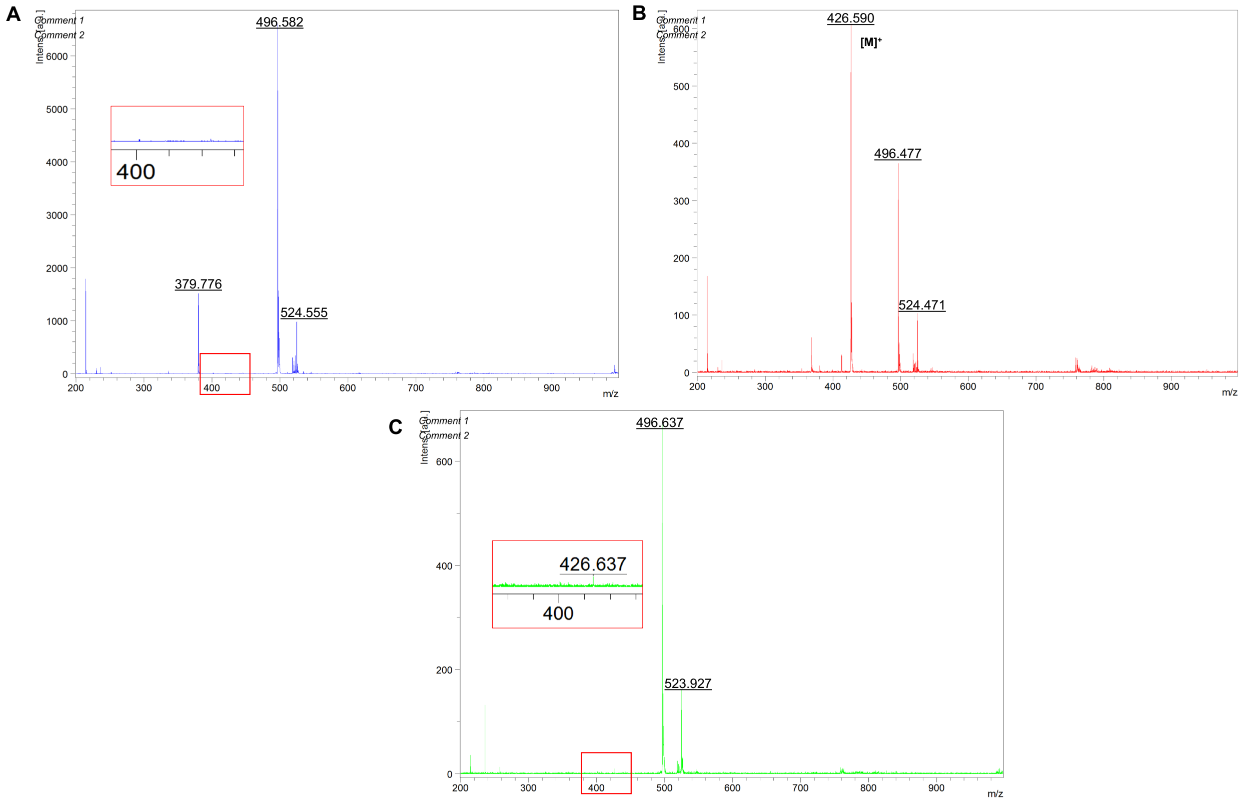


**Fig. S6.** MALDI mass analysis of vehicle (A) and TGR63 treated mice blood serum after 1 h (B) and 24 h (C) of administration. The presence of TGR63 in blood was confirmed from the mass analysis even after 24 h.

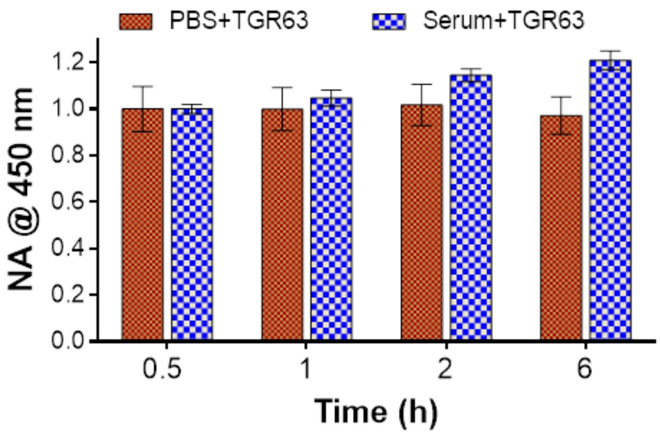


**Fig. S7.** Serum stability of TGR63 under *in vitro* conditions: TGR63 was incubated in PBS (10 mM, pH= 7.4) and blood serum (WT mouse) for different time (0.5, 1, 2 and 6 h) at 37 ºC. Data show the normalized absorbance (NA) of TGR63 at 450 nm recorded at different time intervals, which confirmed the stability of TGR63 in blood serum.

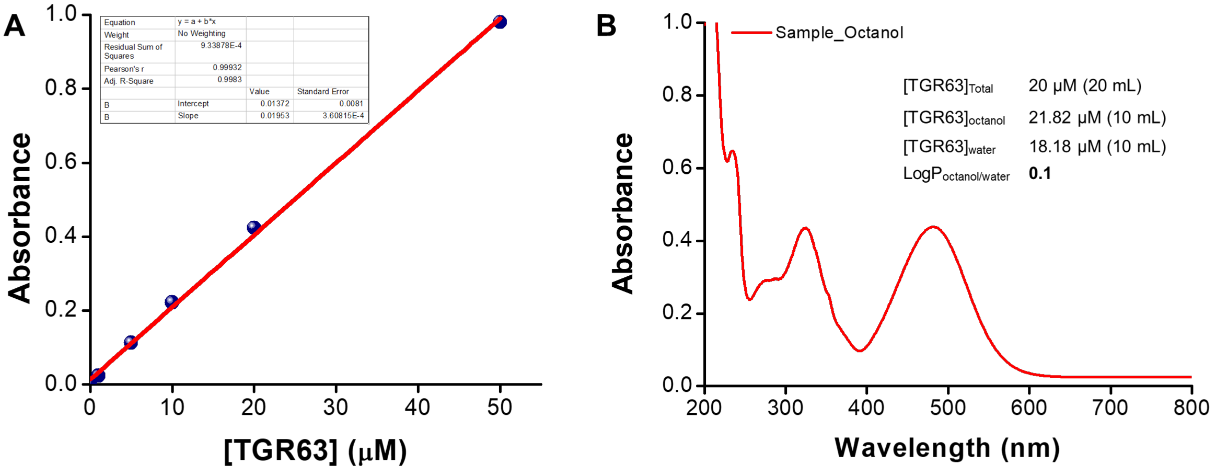


**Fig. S8.** Calculation of LogP. (A) Standard concentration curve obtained by measuring absorbance at 480 nm for 1, 5, 10, 20 and 50 μM of TGR63 in octanol. (B) Absorbance of octanol layer (Sample_Octanol) and calculation of LogP.

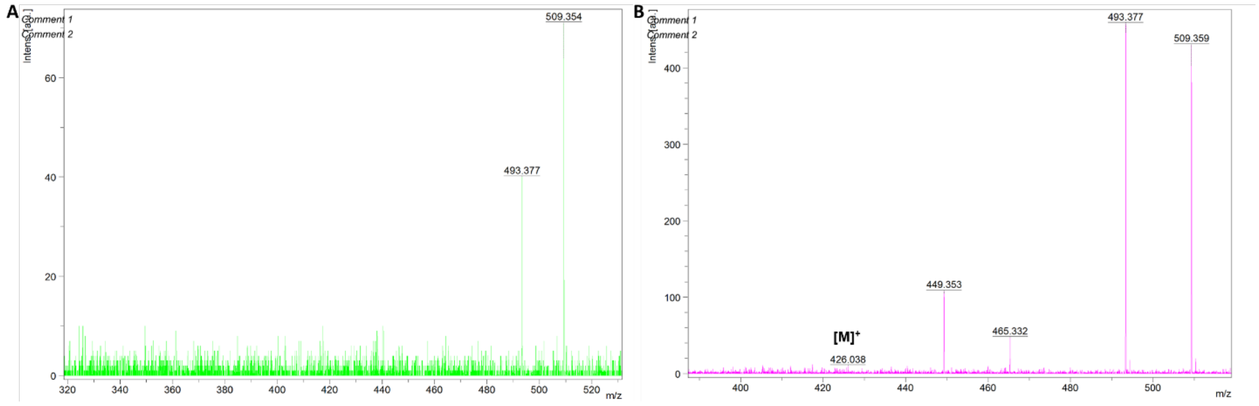


**Fig. S9.** MALDI mass analysis of vehicle (A) and TGR63 (B) treated mouse brain lysate after 1 h. The absence of any characteristic mass peaks in vehicle treated control sample confirm the presence of TGR63 in treated mice brain.

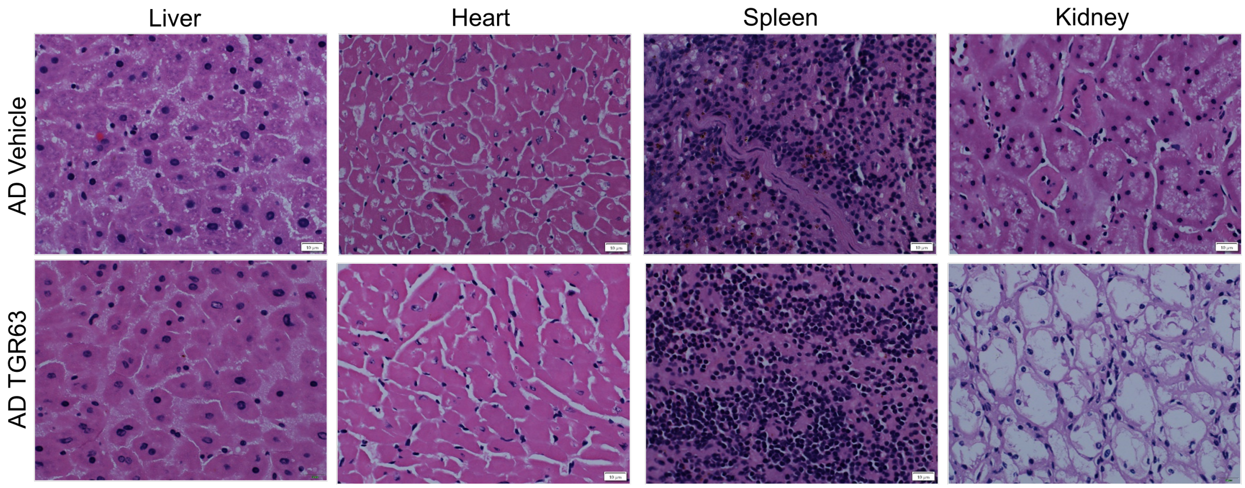


**Fig. S10.** Evaluation of organ toxicity of TGR63. Bright field images of vehicle and TGR63 treated APP/PS1 mice organs (liver, heart, spleen and kidney) stained with hematoxylin and eosin. TGR63 treated mouse organs showed the healthy nature like vehicle treated control and confirmed the biocompatibility and nontoxic nature of TGR63. Scale bar: 10 µm.

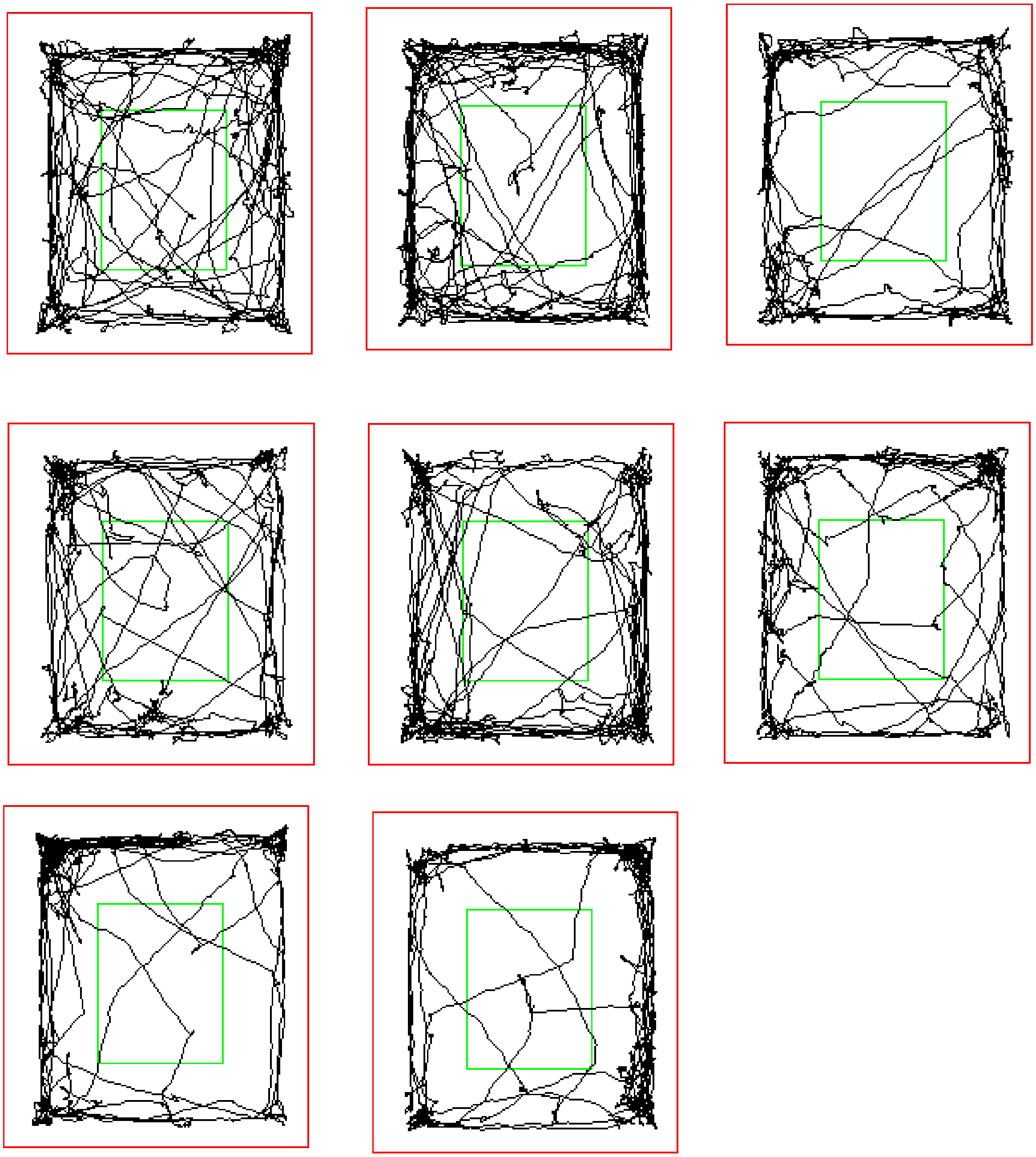

**Fig. S11.** The locomotion of vehicle treated WT mice cohort during OF test.

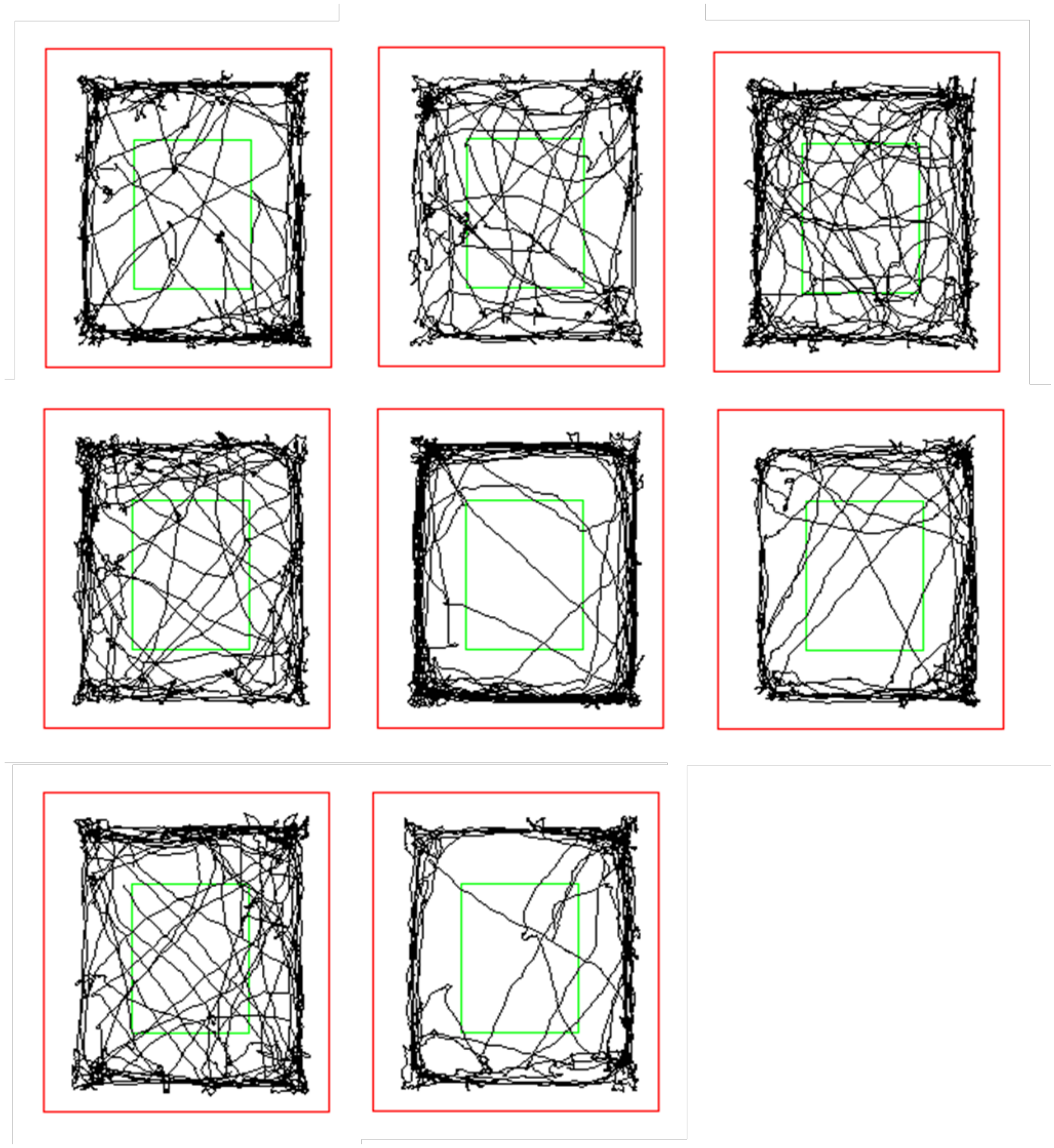

**Fig. S12.** The locomotion of TGR63 treated WT mice cohort during OF test.

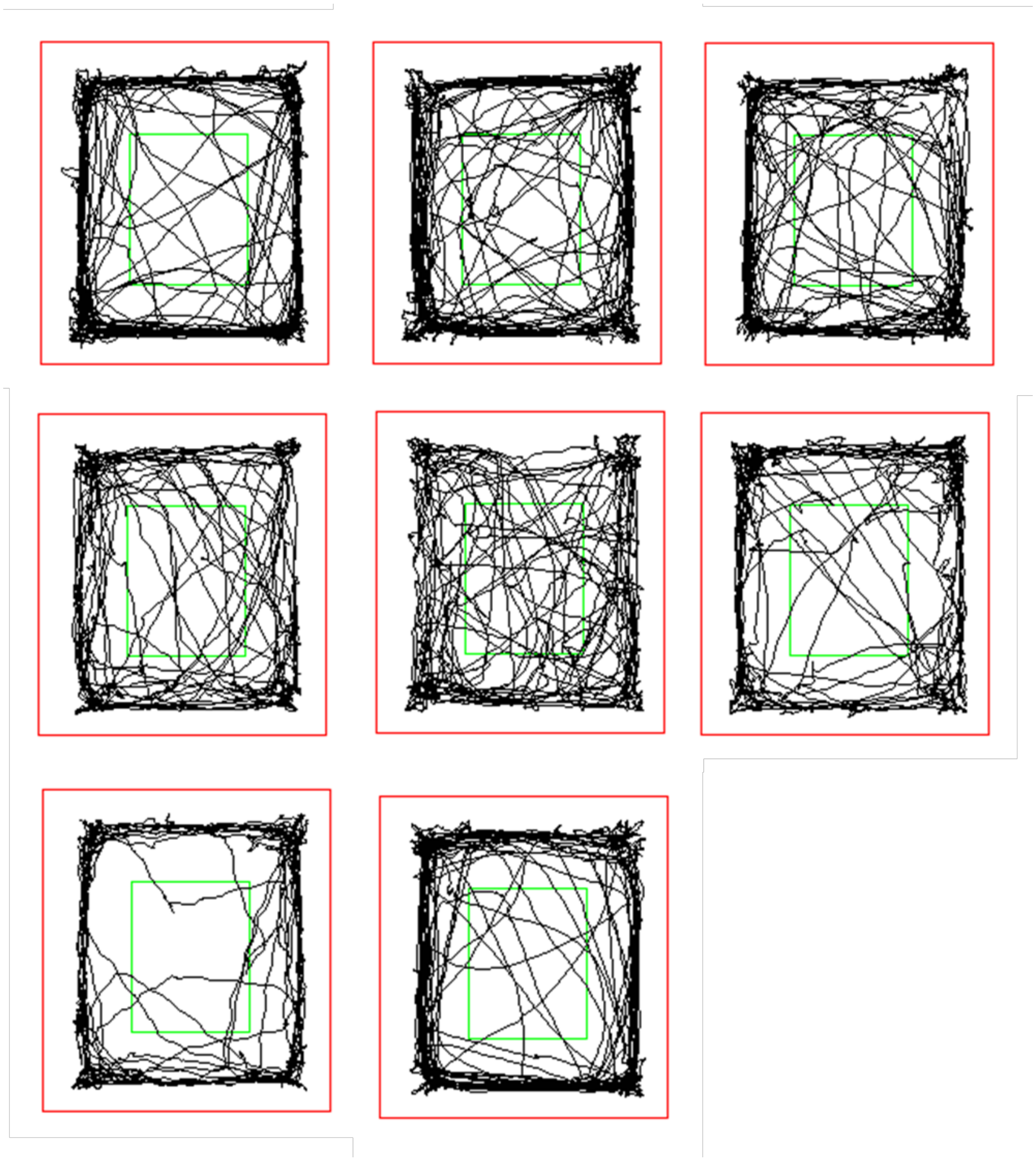

**Fig. S13.** The locomotion of vehicle treated AD mice cohort during OF test.

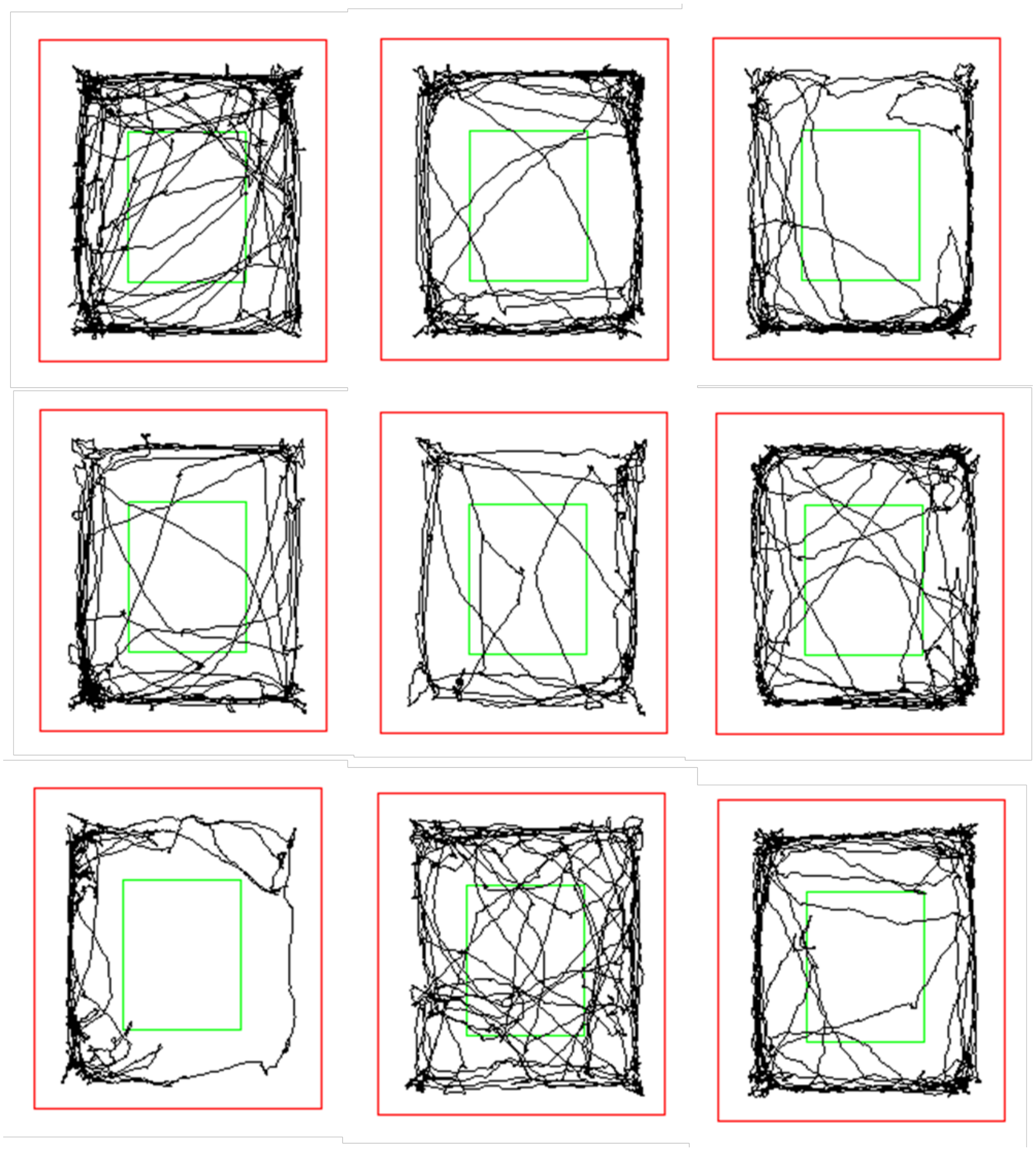

**Fig. S14.** The locomotion of TGR63 treated AD mice cohort during OF test.

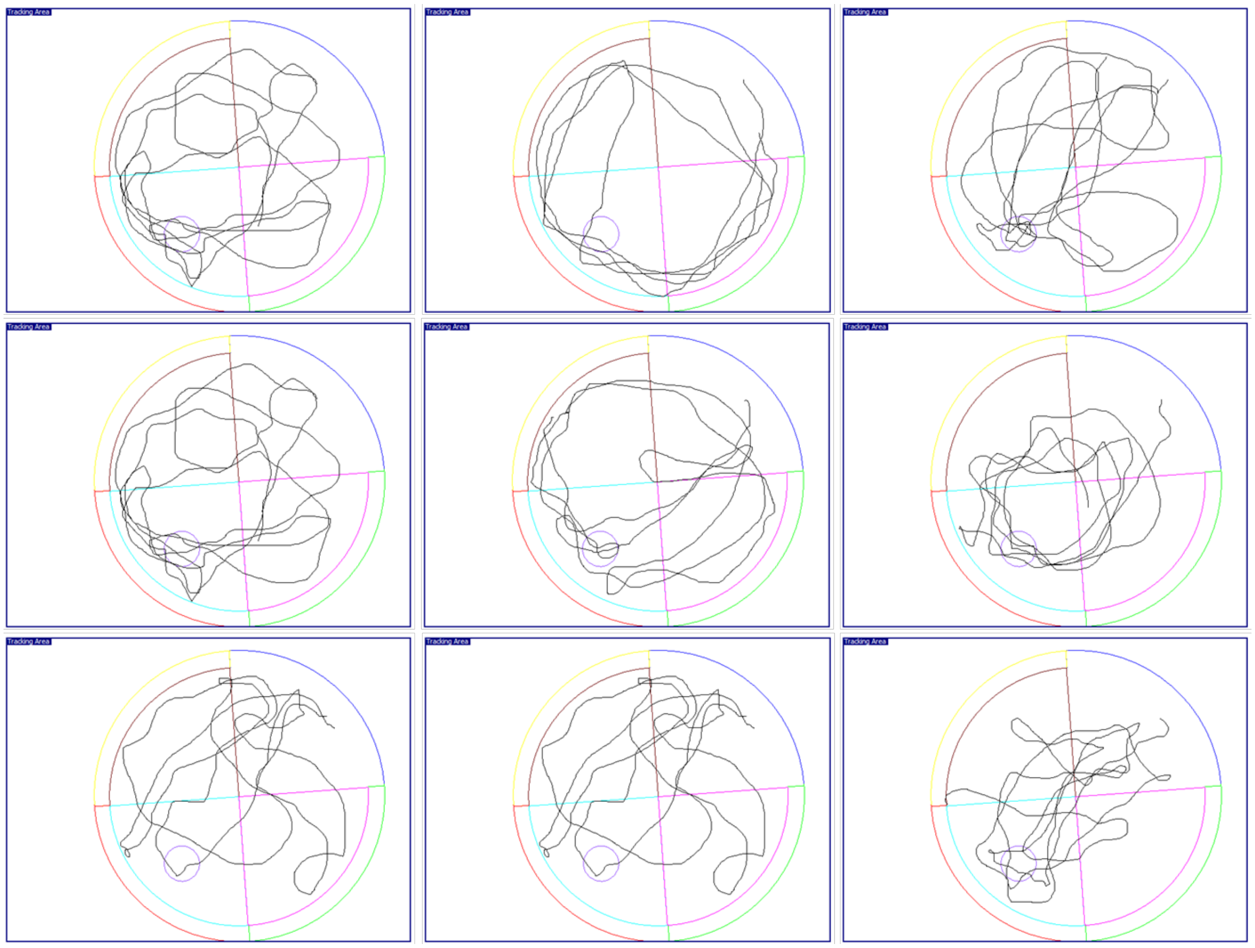

**Fig. S15.** The trajectory of vehicle treated WT mice cohort during MWM probe trail (without platform).

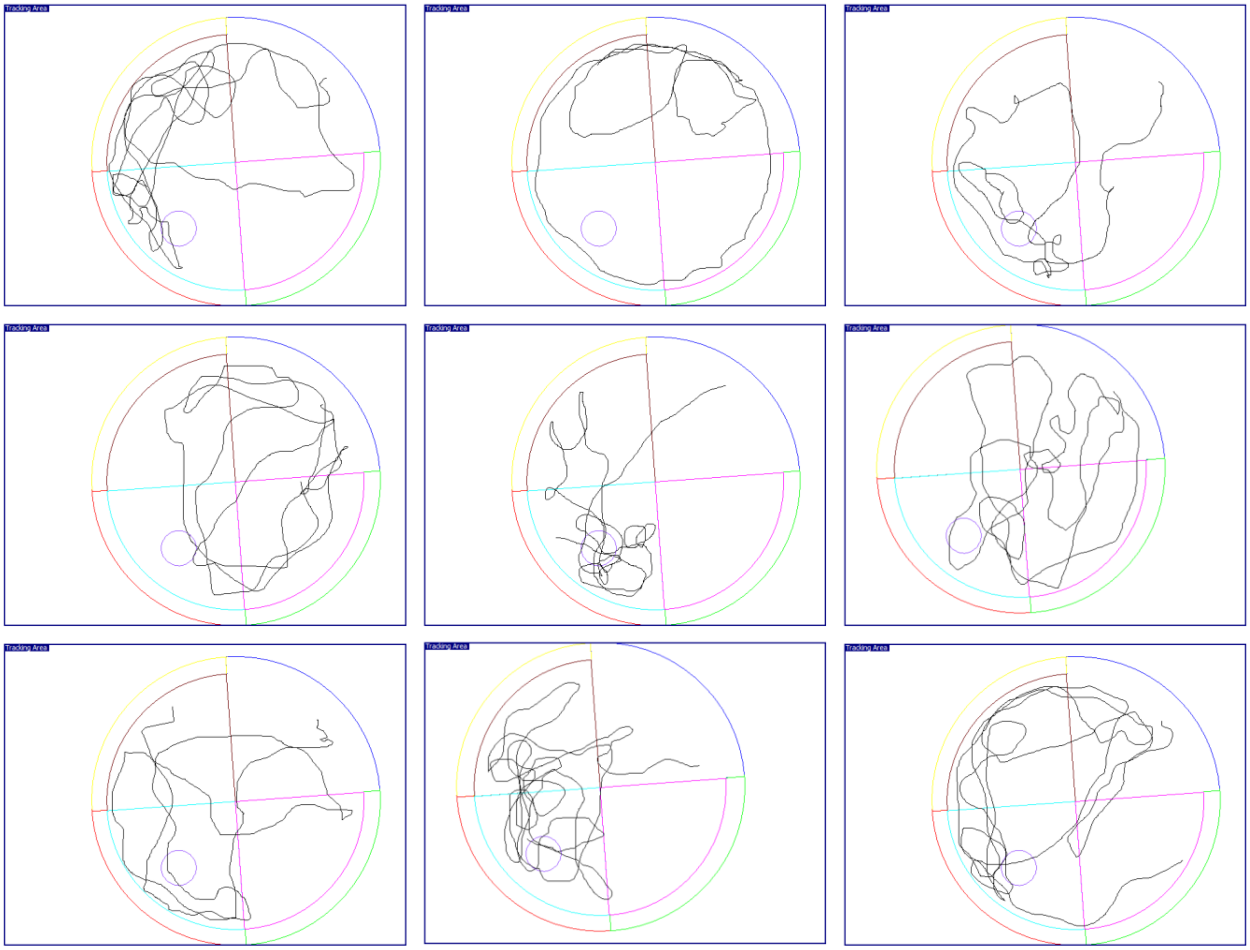


**Fig. S16.** The trajectory of TGR63 treated WT mice cohort during MWM probe trail (without platform).

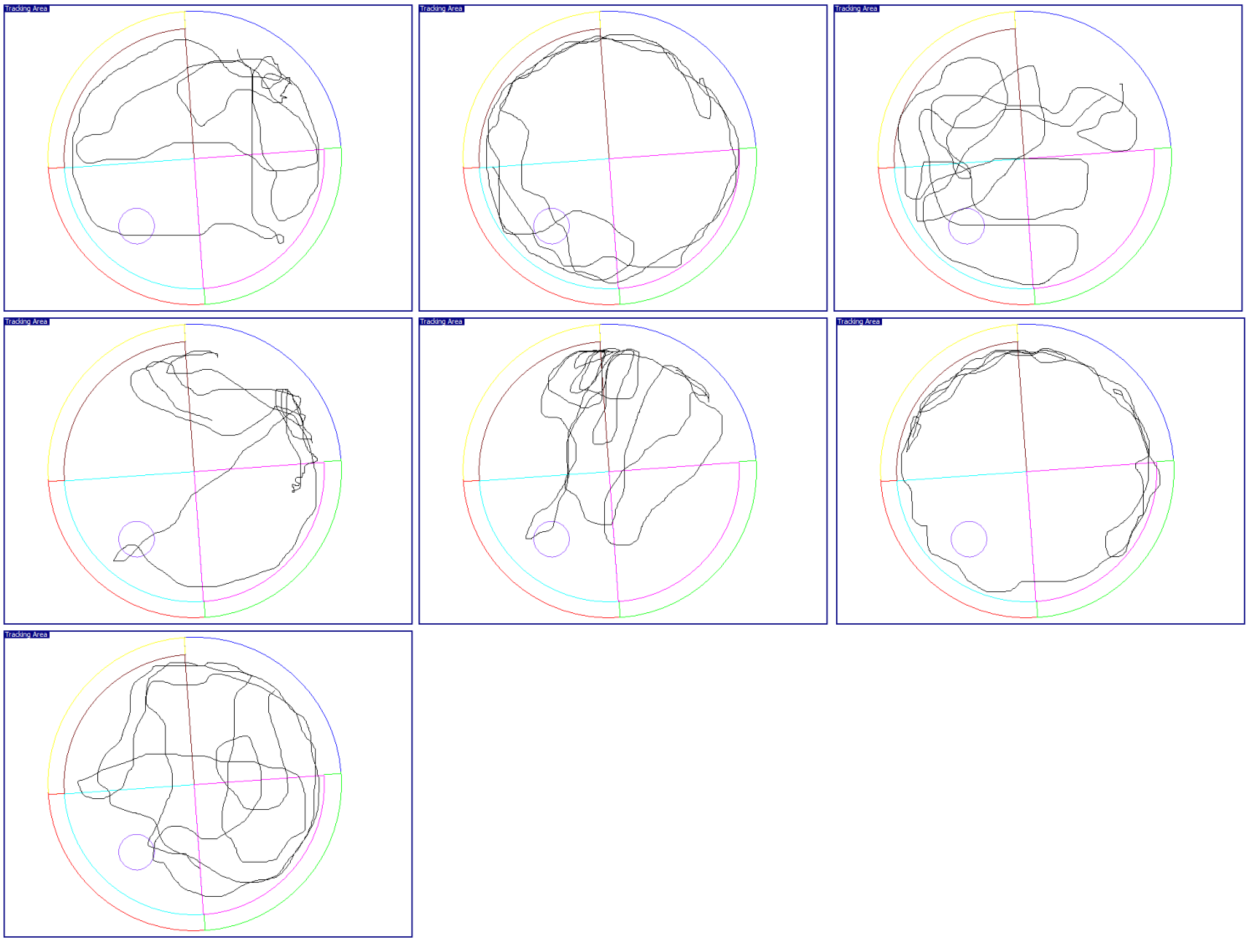

**Fig. S17.** The trajectory of vehicle treated AD mice cohort during MWM probe trail (without platform).

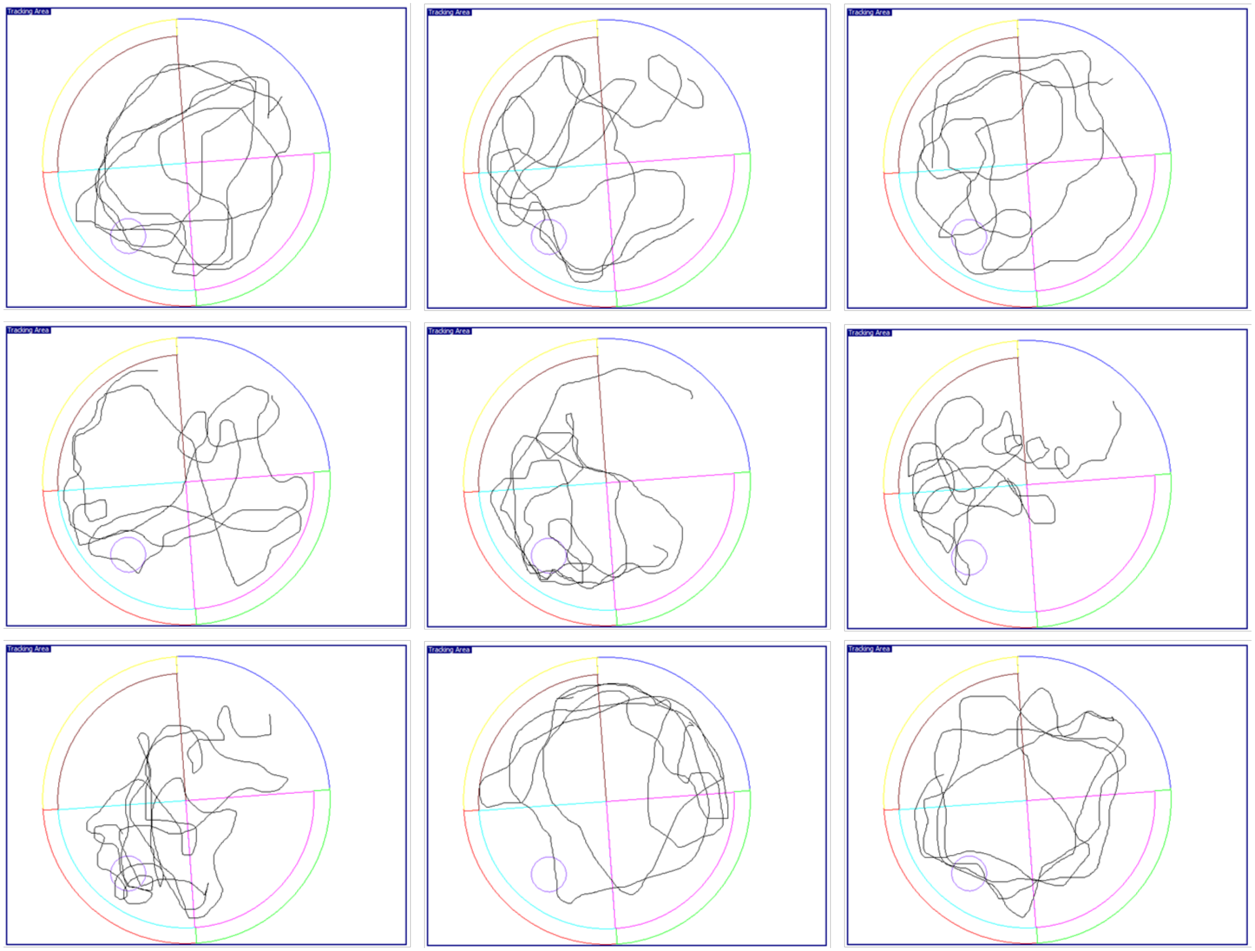


**Fig. S18.** The trajectory of TGR63 treated AD mice cohort during MWM probe trail (without platform).

**Data file S1.** Characterization of data of TGR60-65.

$^1$H NMR

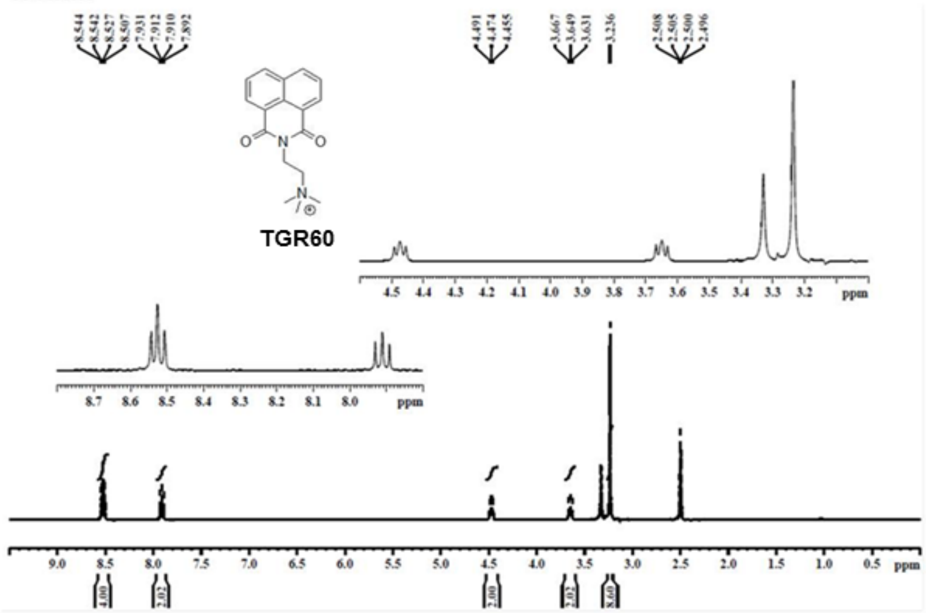


$^{13}$C NMR

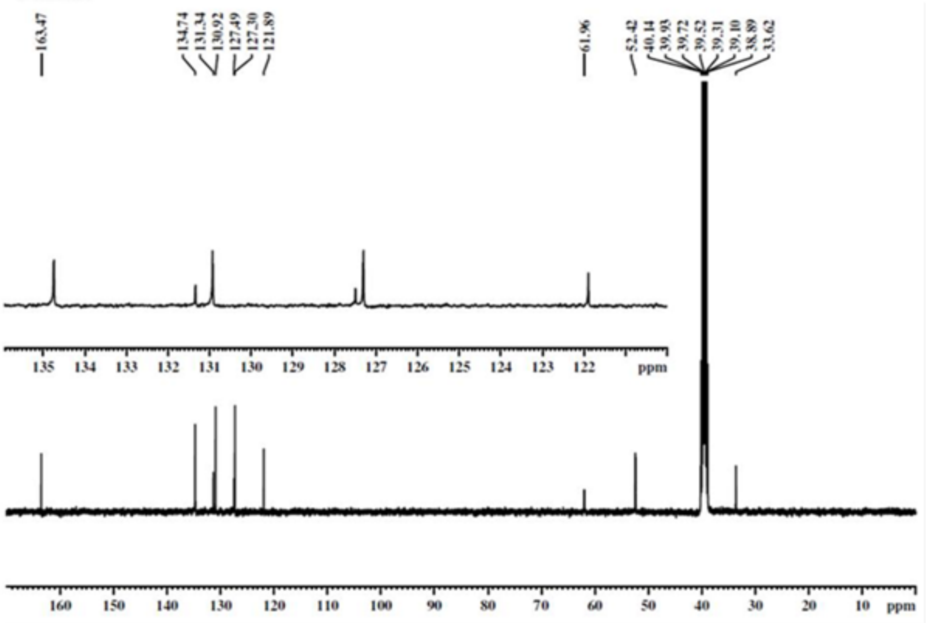


HRMS

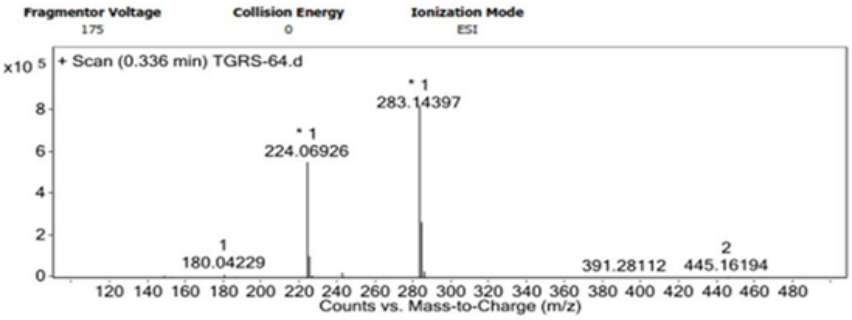

### $^{1}$H NMR

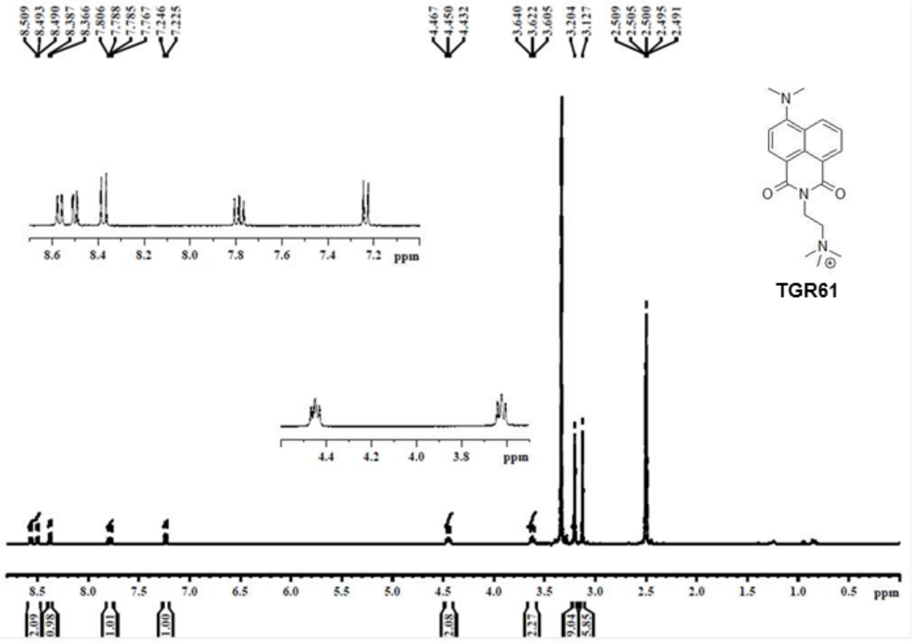


### $^{13}$C NMR

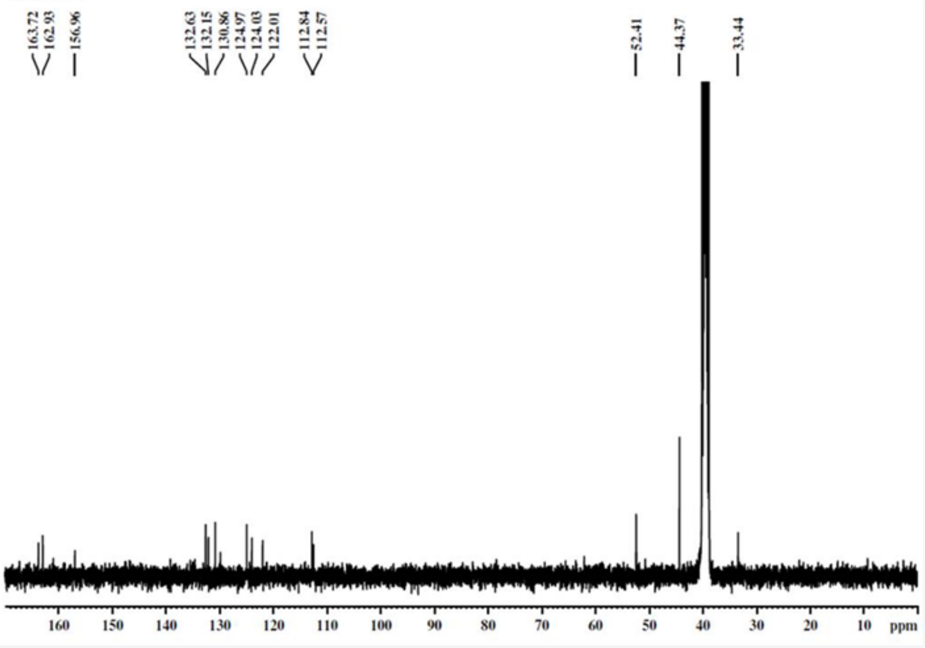


### HRMS

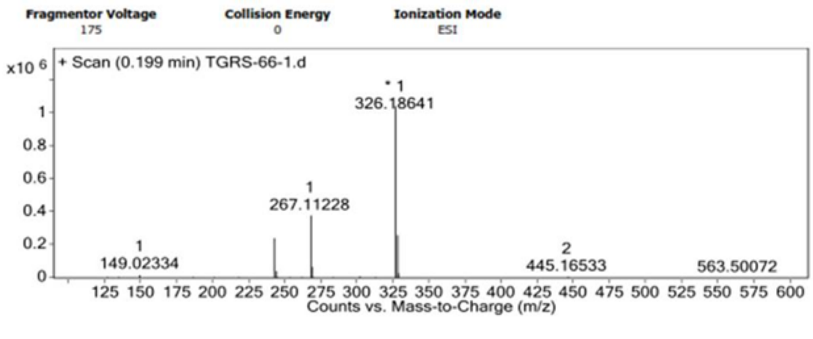

**$^1$H NMR**

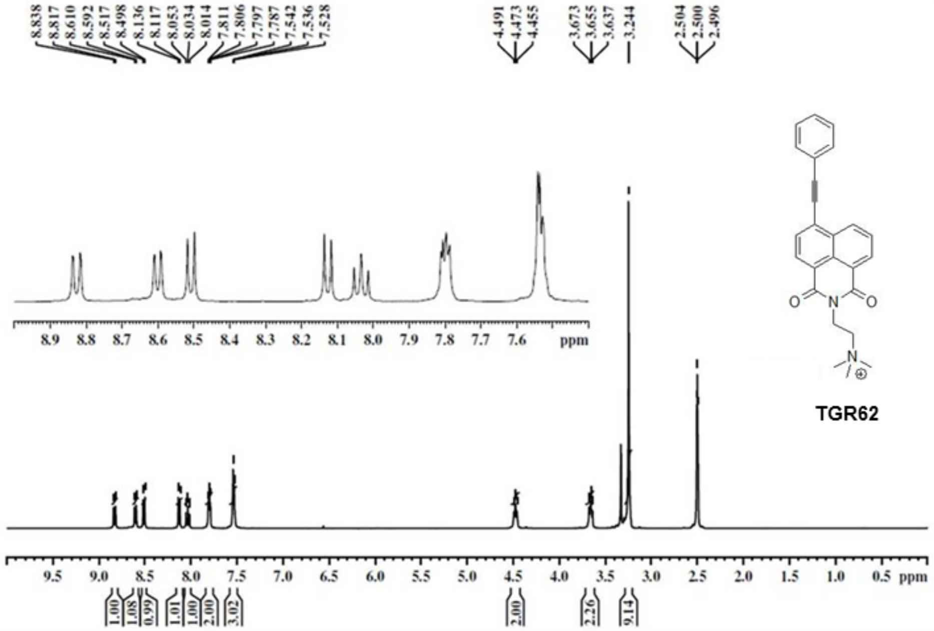


**$^{13}$C NMR**

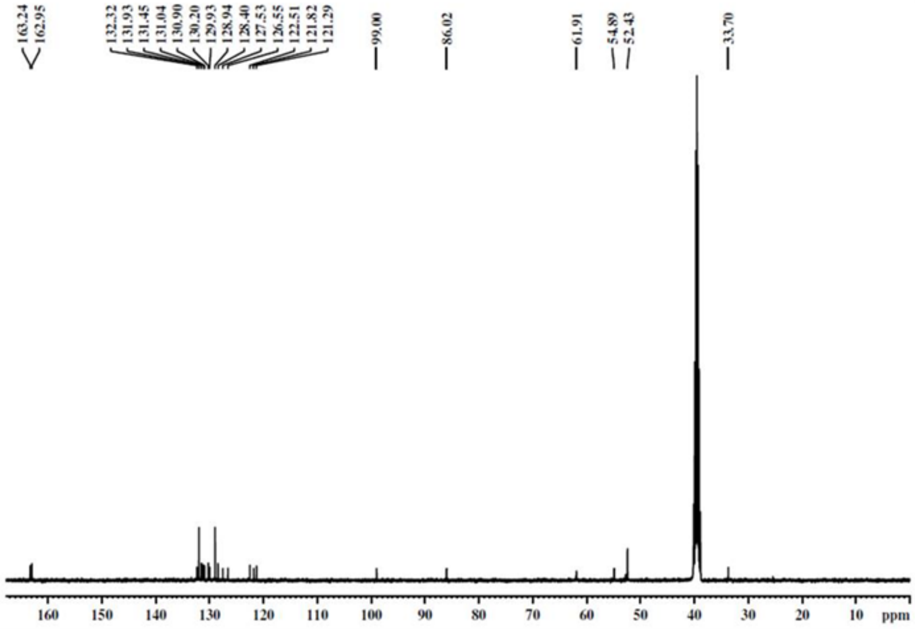


**HRMS**

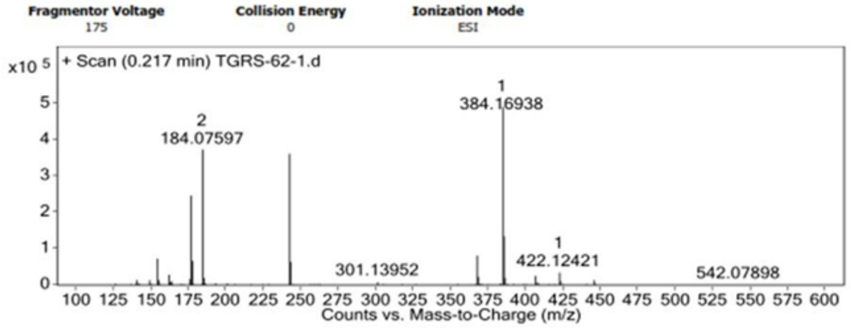

**[1]H NMR**

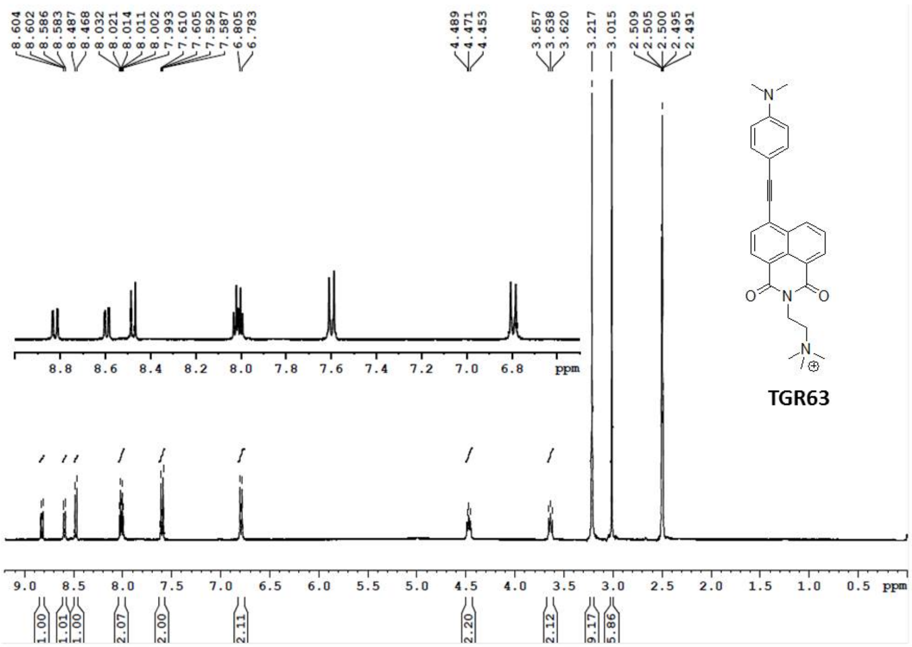


**[13]C NMR**

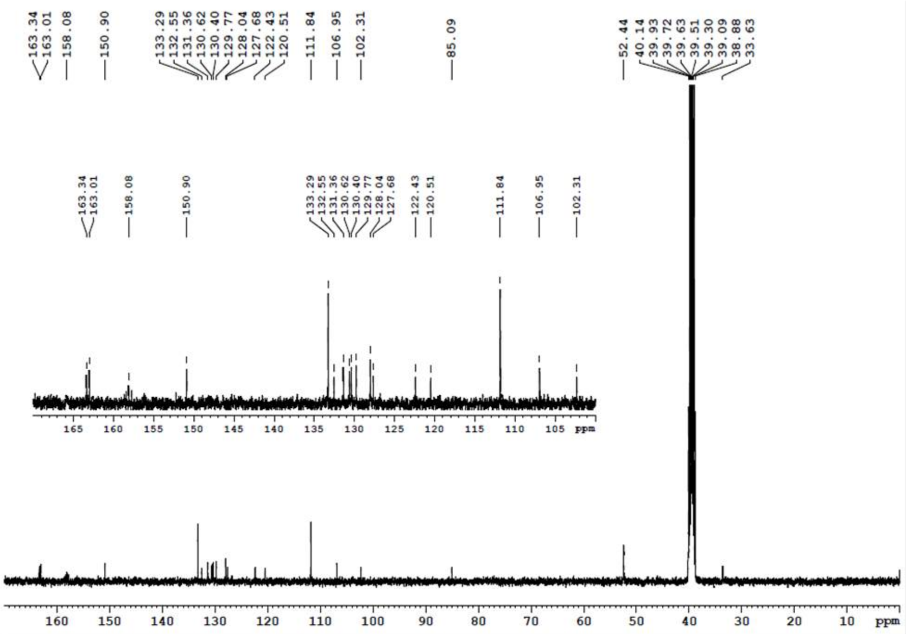


**HRMS**

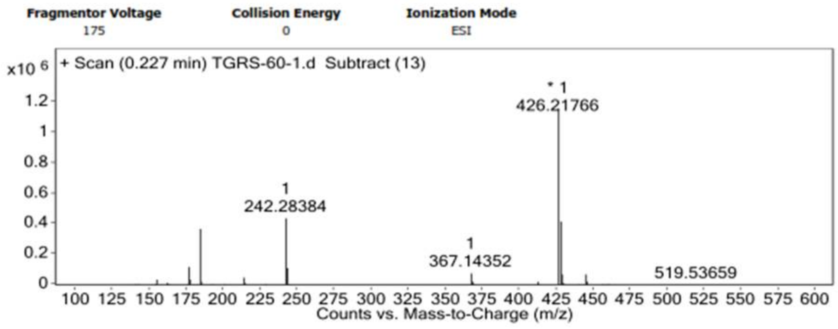

### $^1$H NMR

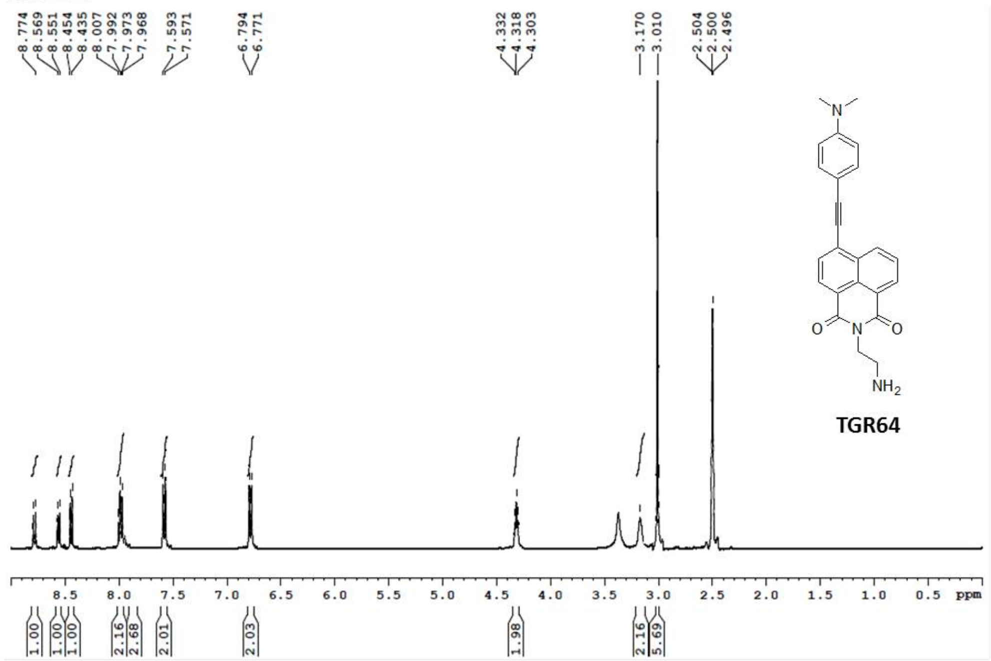


### $^{13}$C NMR

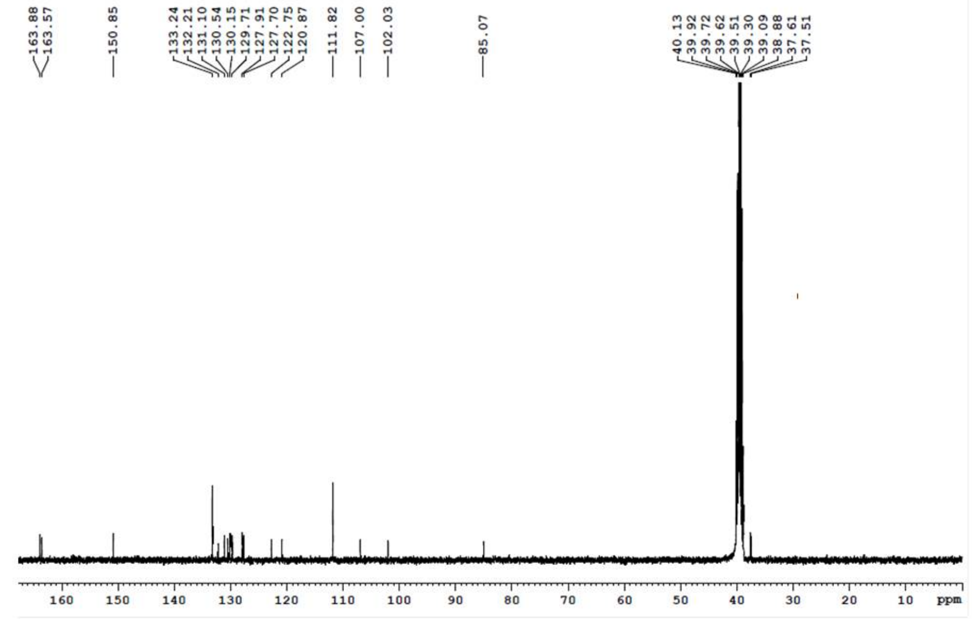


### HRMS

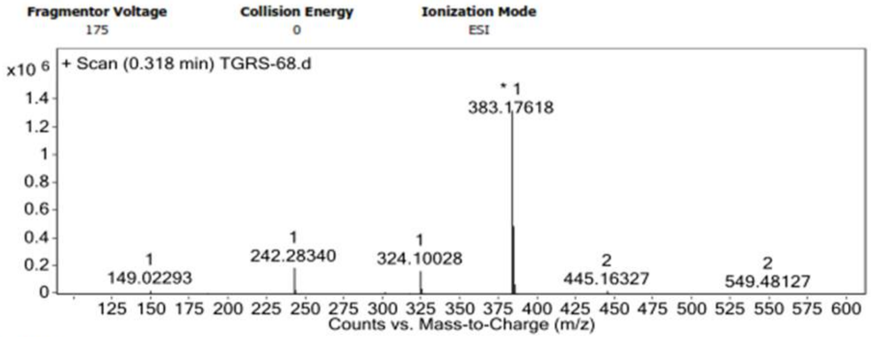

$^{1}H$ NMR

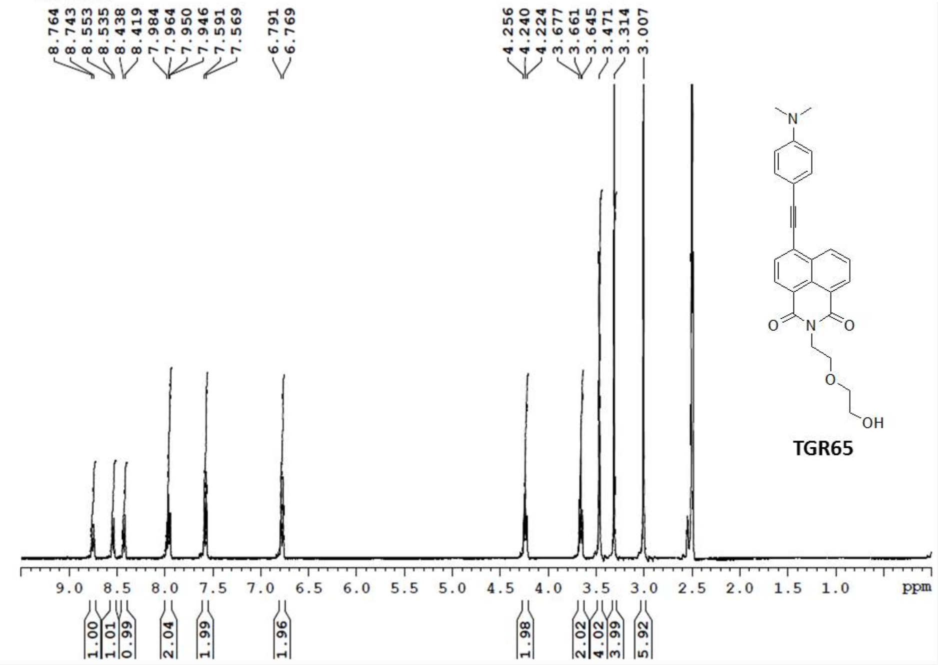


$^{13}C$ NMR

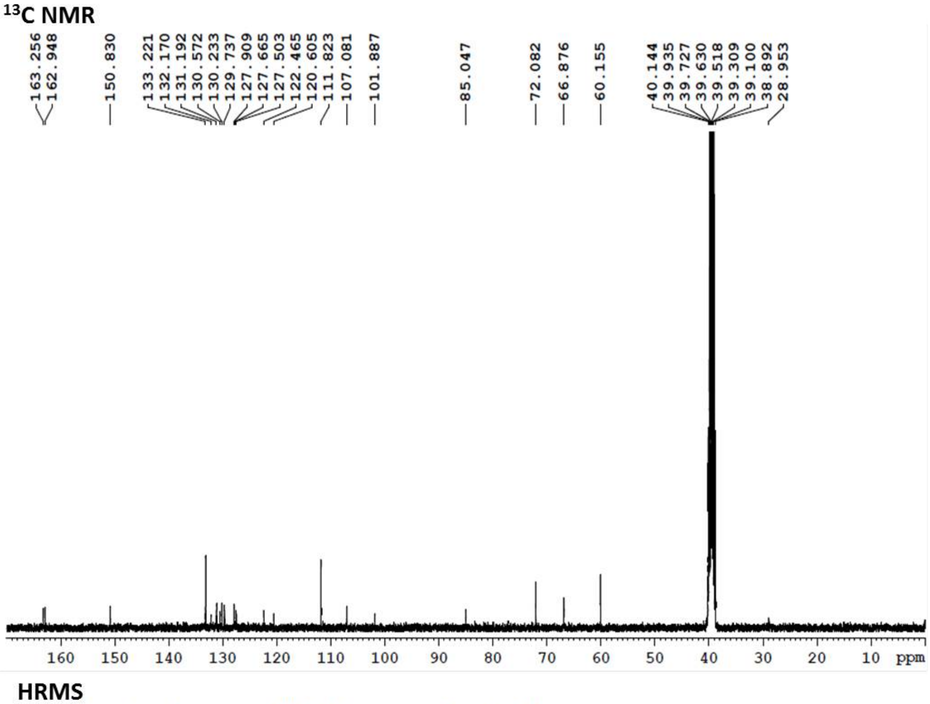


HRMS

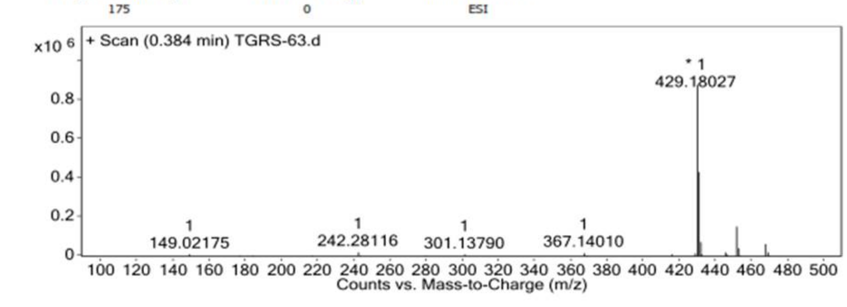

### LCMS characterization of TGR63:

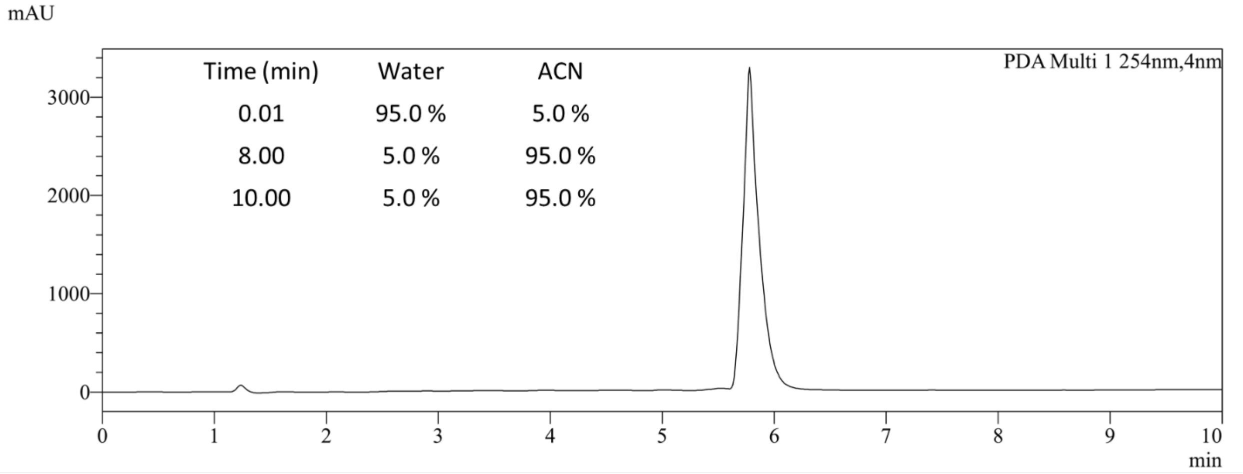


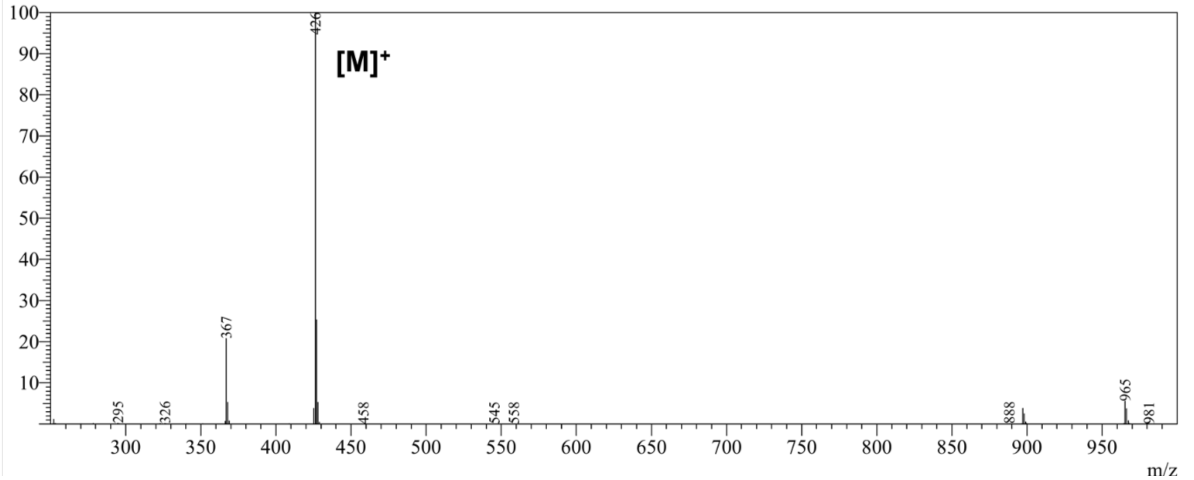

**Statically analysis of Fig. 5B**

| 2way ANOVA Tabular results | A Data Set-A | B Data Set-B | C Data Set-C | D Data Set-D | E Data Set-E |
|---|---|---|---|---|---|
| | Y | Y | Y | Y | Y |
| 1 Table Analyzed | Data 1 | | | | |
| 2 | | | | | |
| 3 Two-way ANOVA | Ordinary | | | | |
| 4 Alpha | 0.05 | | | | |
| 5 | | | | | |
| 6 Source of Variation | % of total variation | P value | P value summary | Significant? | |
| 7 Interaction | 51.69 | < 0.0001 | **** | Yes | |
| 8 Treatment | 8.747 | 0.0087 | ** | Yes | |
| 9 Genotype | 7.831 | 0.0125 | * | Yes | |
| 10 | | | | | |
| 11 ANOVA table | SS | DF | MS | F (DFn, DFd) | P value |
| 12 Interaction | 5.780e+006 | 1 | 5.780e+006 | F (1, 29) = 46.79 | P < 0.0001 |
| 13 Treatment | 978105 | 1 | 978105 | F (1, 29) = 7.918 | P = 0.0087 |
| 14 Genotype | 875730 | 1 | 875730 | F (1, 29) = 7.089 | P = 0.0125 |
| 15 Residual | 3.583e+006 | 29 | 123535 | | |

**Statically analysis of Fig. 5C**

| 2way ANOVA Tabular results | A Data Set-A | B Data Set-B | C Data Set-C | D Data Set-D | E Data Set-E |
|---|---|---|---|---|---|
| | Y | Y | Y | Y | Y |
| 1 Table Analyzed | Data 1 | | | | |
| 2 | | | | | |
| 3 Two-way ANOVA | Ordinary | | | | |
| 4 Alpha | 0.05 | | | | |
| 5 | | | | | |
| 6 Source of Variation | % of total variation | P value | P value summary | Significant? | |
| 7 Interaction | 25.83 | 0.0016 | ** | Yes | |
| 8 Treatment | 2.623 | 0.2810 | ns | No | |
| 9 Group | 1.756 | 0.3763 | ns | No | |
| 10 | | | | | |
| 11 ANOVA table | SS | DF | MS | F (DFn, DFd) | P value |
| 12 Interaction | 662.0 | 1 | 662.0 | F (1, 32) = 11.84 | P = 0.0016 |
| 13 Treatment | 67.22 | 1 | 67.22 | F (1, 32) = 1.203 | P = 0.2810 |
| 14 Group | 45.00 | 1 | 45.00 | F (1, 32) = 0.8050 | P = 0.3763 |
| 15 Residual | 1789 | 32 | 55.90 | | |

**Statically analysis of Fig. 5D**

| 2way ANOVA Tabular results | A | B | C | D | E |
|---|---|---|---|---|---|
| | Data Set-A | Data Set-B | Data Set-C | Data Set-D | Data Set-E |
| | Y | Y | Y | Y | Y |
| 1 Table Analyzed | Data 1 | | | | |
| 2 | | | | | |
| 3 Two-way ANOVA | Ordinary | | | | |
| 4 Alpha | 0.05 | | | | |
| 5 | | | | | |
| 6 Source of Variation | % of total variation | P value | P value summary | Significant? | |
| 7 Interaction | 31.50 | 0.0004 | *** | Yes | |
| 8 Row Factor | 2.167 | 0.3091 | ns | No | |
| 9 Column Factor | 1.430 | 0.4074 | ns | No | |
| 10 | | | | | |
| 11 ANOVA table | SS | DF | MS | F (DFn, DFd) | P value |
| 12 Interaction | 117065 | 1 | 117065 | F (1, 32) = 15.53 | P = 0.0004 |
| 13 Row Factor | 8053 | 1 | 8053 | F (1, 32) = 1.068 | P = 0.3091 |
| 14 Column Factor | 5314 | 1 | 5314 | F (1, 32) = 0.7049 | P = 0.4074 |
| 15 Residual | 241219 | 32 | 7538 | | |

**Statically analysis of Fig. 5F**

| 2way ANOVA Tabular results | A | B | C | D | E |
|---|---|---|---|---|---|
| | Data Set-A | Data Set-B | Data Set-C | Data Set-D | Data Set-E |
| | Y | Y | Y | Y | Y |
| 1 Table Analyzed | Data 1 | | | | |
| 2 | | | | | |
| 3 Two-way ANOVA | Ordinary | | | | |
| 4 Alpha | 0.05 | | | | |
| 5 | | | | | |
| 6 Source of Variation | % of total variation | P value | P value summary | Significant? | |
| 7 Interaction | 21.66 | < 0.0001 | **** | Yes | |
| 8 Treatment | 22.53 | < 0.0001 | **** | Yes | |
| 9 Group | 36.85 | < 0.0001 | **** | Yes | |
| 10 | | | | | |
| 11 ANOVA table | SS | DF | MS | F (DFn, DFd) | P value |
| 12 Interaction | 4509 | 1 | 4509 | F (1, 30) = 29.34 | P < 0.0001 |
| 13 Treatment | 4690 | 1 | 4690 | F (1, 30) = 30.52 | P < 0.0001 |
| 14 Group | 7671 | 1 | 7671 | F (1, 30) = 49.92 | P < 0.0001 |
| 15 Residual | 4610 | 30 | 153.7 | | |

**Statically analysis of Fig. 5G**

| 2way ANOVA Tabular results | A Data Set-A Y | B Data Set-B Y | C Data Set-C Y | D Data Set-D Y | E Data Set-E Y |
|---|---|---|---|---|---|
| 1 Table Analyzed | Data 1 | | | | |
| 2 | | | | | |
| 3 Two-way ANOVA | Ordinary | | | | |
| 4 Alpha | 0.05 | | | | |
| 5 | | | | | |
| 6 Source of Variation | % of total variation | P value | P value summary | Significant? | |
| 7 Interaction | 14.67 | 0.0028 | ** | Yes | |
| 8 Treatment | 16.94 | 0.0015 | ** | Yes | |
| 9 Group | 29.43 | < 0.0001 | **** | Yes | |
| 10 | | | | | |
| 11 ANOVA table | SS | DF | MS | F (DFn, DFd) | P value |
| 12 Interaction | 3672 | 1 | 3672 | F (1, 29) = 10.67 | P = 0.0028 |
| 13 Treatment | 4240 | 1 | 4240 | F (1, 29) = 12.32 | P = 0.0015 |
| 14 Group | 7368 | 1 | 7368 | F (1, 29) = 21.40 | P < 0.0001 |
| 15 Residual | 9983 | 29 | 344.2 | | |

**Statically analysis of Fig. 5K**

| 2way ANOVA Tabular results | A Data Set-A Y | B Data Set-B Y | C Data Set-C Y | D Data Set-D Y | E Data Set-E Y |
|---|---|---|---|---|---|
| 1 Table Analyzed | Data 1 | | | | |
| 2 | | | | | |
| 3 Two-way ANOVA | Ordinary | | | | |
| 4 Alpha | 0.05 | | | | |
| 5 | | | | | |
| 6 Source of Variation | % of total variation | P value | P value summary | Significant? | |
| 7 Interaction | 17.19 | < 0.0001 | **** | Yes | |
| 8 Quadrant | 64.04 | < 0.0001 | **** | Yes | |
| 9 Group | 8.490e-009 | > 0.9999 | ns | No | |
| 10 | | | | | |
| 11 ANOVA table | SS | DF | MS | F (DFn, DFd) | P value |
| 12 Interaction | 7462 | 3 | 2487 | F (3, 68) = 16.15 | P < 0.0001 |
| 13 Quadrant | 27799 | 1 | 27799 | F (1, 68) = 180.4 | P < 0.0001 |
| 14 Group | 3.686e-006 | 3 | 1.229e-006 | F (3, 68) = 7.974e-009 | P > 0.9999 |
| 15 Residual | 10476 | 68 | 154.1 | | |

**Statically analysis of Fig. 5L**

| 2way ANOVA Tabular results | | A | B | C | D | E |
|---|---|---|---|---|---|---|
| | | Data Set-A | Data Set-B | Data Set-C | Data Set-D | Data Set-E |
| | | Y | Y | Y | Y | Y |
| 1 | Table Analyzed | Data 1 | | | | |
| 2 | | | | | | |
| 3 | Two-way ANOVA | Ordinary | | | | |
| 4 | Alpha | 0.05 | | | | |
| 5 | | | | | | |
| 6 | Source of Variation | % of total variation | P value | P value summary | Significant? | |
| 7 | Interaction | 16.28 | 0.0102 | * | Yes | |
| 8 | Treatment | 5.523 | 0.1220 | ns | No | |
| 9 | Group | 8.323 | 0.0599 | ns | No | |
| 10 | | | | | | |
| 11 | ANOVA table | SS | DF | MS | F (DFn, DFd) | P value |
| 12 | Interaction | 25.78 | 1 | 25.78 | F (1, 33) = 7.427 | P = 0.0102 |
| 13 | Treatment | 8.744 | 1 | 8.744 | F (1, 33) = 2.519 | P = 0.1220 |
| 14 | Group | 13.18 | 1 | 13.18 | F (1, 33) = 3.797 | P = 0.0599 |
| 15 | Residual | 114.5 | 33 | 3.471 | | |
